# GluN2C/D-containing NMDA receptors enhance temporal summation and increase sound-evoked and spontaneous firing in the inferior colliculus

**DOI:** 10.1101/2023.04.27.538607

**Authors:** Audrey C. Drotos, Rachel L. Zarb, Victoria Booth, Michael T. Roberts

## Abstract

Along the ascending auditory pathway, there is a broad shift from temporal coding, which is common in the lower auditory brainstem, to rate coding, which predominates in auditory cortex. This temporal-to-rate transition is particularly prominent in the inferior colliculus (IC), the midbrain hub of the auditory system, but the mechanisms that govern how individual IC neurons integrate information across time remain largely unknown. Here, we report the widespread expression of *Glun2c* and *Glun2d* mRNA in IC neurons. GluN2C/D-containing NMDA receptors are relatively insensitive to voltage-dependent Mg^2+^ block, and thus can conduct current at resting membrane potential. Using in situ hybridization and pharmacology, we show that VIP neurons in the IC express GluN2D-containing NMDA receptors that are activatable by commissural inputs from the contralateral IC. In addition, GluN2C/D-containing receptors have much slower kinetics than other NMDA receptors, and we found that GluN2D-containing receptors facilitate temporal summation of synaptic inputs in VIP neurons. In a model neuron, we show that a GluN2C/D-like conductance interacts with the passive membrane properties of the neuron to alter temporal and rate coding of stimulus trains. Consistent with this, we show in vivo that blocking GluN2C/D-containing receptors decreases both the spontaneous firing rate and the overall firing rate elicited by amplitude-modulated (AM) sounds in many IC neurons. These results suggest that GluN2C/D-containing NMDA receptors influence rate coding for auditory stimuli in the IC by facilitating the temporal integration of synaptic inputs.

**Significance statement:** NMDA receptors are critical components of most glutamatergic circuits in the brain, and the diversity of NMDA receptor subtypes yields receptors with a variety of functions. We found that many neurons in the auditory midbrain express GluN2C and/or GluN2D NMDA receptor subunits, which are less sensitive to Mg^2+^ block than the more commonly expressed GluN2A/B subunits. We showed that GluN2C/D-containing receptors conducted current at resting membrane potential and enhanced temporal summation of synaptic inputs. In a model, we show that GluN2C/D-containing receptors provide additive gain for input-output functions driven by trains of synaptic inputs. In line with this, we found that blocking GluN2C/D-containing NMDA receptors in vivo decreased both spontaneous firing rates and firing evoked by amplitude-modulated sounds.

## Introduction

The inferior colliculus (IC) in the midbrain plays a critical role in auditory processing, functioning as a hub of integration in the auditory pathway (Adams, 1979; Oliver and Huerta, 1992) and as a site where phase-locked ascending inputs are transformed into a firing rate code (Hewitt and Meddis, 1994; Joris et al., 2004). However, the cellular and circuit mechanisms that govern how IC neurons integrate synaptic inputs and convert temporal to rate codes remain largely unknown. In other brain regions, the slow kinetics of NMDA receptors (NMDARs) can play a critical role in temporal integration of synaptic inputs (D’Angelo et al., 1995; Lo et al., 2013). Intriguingly, previous studies have found that many IC neurons express NMDARs that are activated at resting membrane potential (Kelly and Zhang, 2002; Ma et al., 2002; Wu, 2004; Oberle et al., 2022). In line with this previous work, we recently found in adult mice that optogenetic stimulation of commissural inputs elicited excitatory postsynaptic potentials (EPSPs) in IC VIP neurons that had a significant contribution from NMDARs at resting membrane potential (Goyer et al., 2019).

These results suggest that NMDARs that conduct current at resting membrane potential are a common feature of IC circuits, but this is unusual. At most glutamatergic synapses, activation of AMPARs is required to depolarize the membrane potential and remove voltage-dependent Mg^2+^ block of NMDARs, allowing NMDARs to contribute to the response. One potential explanation for this phenomenon is that NMDARs in IC neurons may be less sensitive to Mg^2+^ block due to their subunit composition (Paoletti et al., 2013). NMDARs are heterotetramers comprised of two obligatory GluN1 subunits and a combination of GluN2(A-D) and/or GluN3(A-B) subunits (Traynelis et al., 2010). A residue in the GluN2 subunit confers Mg^2+^ sensitivity (Monyer et al., 1992; Siegler Retchless et al., 2012), with GluN2C/D subunit-containing receptors being much less sensitive to Mg^2+^ block than GluN2A/B subunit-containing receptors. Inclusion of GluN2C/D subunits in NMDARs thus gives rise to receptors with significant currents at resting membrane potential, as has been demonstrated in the mouse barrel cortex (Binshtok et al., 2006).

In addition to their differences in Mg^2+^ sensitivity, NMDARs containing GluN2C or GluN2D subunits have slower kinetics than NMDARs with GluN2A or GluN2B subunits (Paoletti et al., 2013). These slow kinetics can expand the time window for temporal summation of synaptic inputs, as seen in neocortex, where GluN2C/D-containing receptors enable pyramidal neurons to integrate intracortical inputs over broader time windows than GluN2A/B-containing receptors (Kumar and Huguenard, 2003).

Here, we tested whether GluN2C/D-containing NMDARs mediate NMDAR currents in the IC and the role of these receptors in synaptic integration and sound processing. We found that IC VIP neurons, a class of excitatory principal neurons (Goyer et al., 2019), express NMDARs that activate at resting potential. We show that VIP neurons express *Glun2d* mRNA and that many other IC neurons express *Glun2c* and/or *Glun2d* mRNA, pointing to a prominent role for GluN2C/D-containing NMDARs in the IC. NMDAR-mediated responses in VIP neurons were sensitive to GluN2C/D-specific pharmacology and were elicited by commissural inputs from the contralateral IC. Additionally, GluN2C/D-specific antagonists decreased temporal summation of commissural inputs in VIP neurons. In a model neuron, addition of an NMDAR-like conductance lacking Mg^2+^ sensitivity altered temporal and rate coding. In line with this, during in vivo recordings from awake mice, local infusion of a GluN2C/D-specific antagonist decreased spontaneous and sound-evoked firing rates in many IC neurons. Overall, these results provide a molecular mechanism for the presence of NMDAR currents at resting potential in many IC neurons and show that the widespread expression of GluN2C/D subunits in the IC influences sound processing.

## Materials and Methods

### Animals

All experiments were approved by the University of Michigan Institutional Animal Care and Use Committee and were in accordance with NIH guidelines for the care and use of laboratory animals. Animals were kept on a 12-hour day/night cycle with ad libitum access to food and water. To visualize VIP neurons, VIP-IRES-Cre mice (*Vip^TM1(cre)Zjh^*/J, Jackson Laboratory, stock #010908) (Taniguchi et al., 2011) were crossed with Ai14 reporter mice (B6.Cg-*Gt(ROSA)26Sor^TM^ ^14(CAG–tdTomato)Hze^*/J, Jackson Laboratory, stock #007914) (Madisen et al., 2010) so that the fluorescent protein tdTomato was expressed in VIP neurons (Beebe et al., 2022). Since mice on the C57BL/6J background undergo age-related hearing loss after 3 months of age (Zheng et al., 1999), all experiments were performed on mice between postnatal days (P)30 – 66.

### Brain slice preparation

Whole-cell patch-clamp recordings were performed in acutely prepared IC slices from VIP-IRES-Cre x Ai14 mice. Male (n = 18) and female (n = 15) mice aged P30 – P66 were used. Mice were deeply anesthetized with isoflurane and then rapidly decapitated. The brain was removed and the IC was dissected in 34 °C artificial cerebrospinal (ACSF) solution containing (in mM): 125 NaCl, 12.5 glucose, 25 NaHCO_3_, 3 KCl, 1.25 NaH_2_PO_4_, 1.5 CaCl_2_, 1 MgSO_4_, 3 sodium pyruvate and 0.40 L-ascorbic acid (Acros Organics), bubbled with 5% CO_2_ in 95% O_2_. 200 µm coronal slices containing the IC were cut using a vibrating microtome (VT1200S, Leica Biosystems). Slices were incubated at 34 °C for 30 min in ACSF bubbled with 5% CO_2_ in 95% O_2_ and then placed at room temperature for at least 30 min before initiating recordings. Recordings were targeted to tdTomato-expressing VIP neurons in the central nucleus of the IC using a Nikon FN1 or Olympus BX51 microscope. All chemicals were obtained from Thermo Fisher Scientific unless otherwise noted.

### Voltage-clamp electrophysiological recordings

Slices were placed in a recording chamber and continuously perfused at a rate of ∼2 ml/min with 34 °C ACSF bubbled in 5% CO_2_/95% O_2_. Whole-cell voltage-clamp recordings were performed with an Axopatch 200A patch clamp amplifier (Axon Instruments). For each recording, series resistance compensation was performed using 80% prediction and 80% correction, and whole cell capacitance was compensated. Series resistance for all neurons included in this study was <15 MΩ. Data were low-pass filtered at 10 kHz, sampled at 50 kHz with a National Instruments PCIe-6343 data acquisition board, and acquired using custom software written in Igor Pro (Sutter Instrument). Recording pipettes were pulled from borosilicate glass capillaries (outer diameter 1.5 mm, inner diameter 0.86 mm, Sutter Instrument) with a P-1000 microelectrode puller (Sutter Instrument) and filled with an internal solution containing (in mM): 115 CsOH, 115 D-gluconic acid, 7.76 CsCl, 0.5 EGTA, 10 HEPES, 10 Na_2_ phosphocreatine, 4 MgATP, 0.3 NaGTP, supplemented with 0.1% biocytin (w/v), pH adjusted to 7.4 with CsOH and osmolality to 290 mmol/kg with sucrose. Voltage clamp recordings were not corrected for the liquid junction potential. All electrophysiological recordings were targeted to neurons in the central nucleus of the IC, but it is possible a small number of recordings were performed in the dorsal cortex of the IC.

To apply glutamate puffs to brain slices, we pulled puffer pipettes from borosilicate glass (outer diameter 1.5 mm, inner diameter 0.86 mm, Sutter Instrument) to a resistance of 3.5 – 5.0 MΩ using a P-1000 microelectrode puller and filled them with 300 μM glutamate (Sigma, # G5889) dissolved in a vehicle solution containing (in mM): 125 NaCl, 3 KCl, 12.5 glucose and 3 HEPES. The solution was balanced to a pH of 7.40 with NaOH. Puffer pipettes were connected to a pressure ejection system built based on the OpenSpritzer design (Forman et al., 2017). The tips of puffer pipettes were placed near the soma of the recorded cell, and 10 ms puffs were presented either 10 ms or 30 ms apart (30 ms for drug conditions) with 5 puffs presented per condition. Puffs containing only the vehicle solution did not elicit any response in the neurons tested (data not shown).

To examine whether NMDAR activation in VIP neurons at resting membrane potential is mediated by NMDARs containing GluN2C/D subunits, we performed whole-cell patch-clamp recordings targeted to VIP neurons as described above. To isolate the contribution of NMDARs, recordings were performed in the presence of 10 µM NBQX, an AMPAR antagonist. Glutamate was puffed onto VIP neurons during bath application of 1.5 μM PPDA (Hello Bio, #HB0531), a GluN2C/D specific antagonist (Lozovaya et al., 2004; Li et al., 2020; Jing et al., 2022), or 20 μM CIQ (Hello Bio, HB0197), a GluN2C/D specific positive allosteric modulator (Mullasseril et al., 2010; Feng et al., 2014; Zhang et al., 2014; Nouhi et al., 2018; Liu et al., 2021). The concentrations of PPDA and CIQ used throughout this study are based on previous studies that used PPDA (Lozovaya et al., 2004; Li et al., 2020; Jing et al., 2022) or CIQ (Feng et al., 2014, 2014; Nouhi et al., 2018; Liu et al., 2021). All drugs were washed in for 10 minutes before testing how the drugs affected responses to glutamate puffs.

CIQ and PPDA were prepared in 0.04% and 0.003% DMSO (volume/volume), respectively. To test for vehicle effects, we performed control experiments where puff-elicited EPSCs were compared between control and DMSO (vehicle) conditions. No differences in puff-elicited EPSCs were found after DMSO wash-in (Linear mixed model tests: amplitude, β = -5.96, *p* = 0.42, CI [-20.2, 8.28], halfwidth, β =7.50, *p* = 0.36, CI [-8.36, 23.37], rise time, β = 13.87, *p* = 0.53, CI [-30.16, 57.91], decay tau, β = -115.62, *p* = 0.20, CI [-281.84, 50.60]).

### Mg^2+^-free voltage-clamp recordings

To determine what proportion of NMDARs on VIP neurons contain GluN2C/D subunits vs GluN2A/B subunits, we performed whole-cell patch-clamp recordings targeted to VIP neurons as described above, except immediately prior to patching brain slices were transferred to a Mg^2+^-free ACSF solution containing (in mM): 125 NaCl, 12.5 glucose, 25 NaHCO_3_, 3 KCl, 1.25 NaH_2_PO_4_, 2.5 CaCl_2_, 0 MgSO_4_, 3 sodium pyruvate and 0.40 L-ascorbic acid (Acros Organics), bubbled with 5% CO_2_ in 95% O_2_. To isolate the contribution of NMDARs, recordings were performed in the presence of 10 µM NBQX. To determine whether GluN2A/B-containing NMDARs were present on VIP neurons, we first performed control glutamate puffs and then puffs after bath application of 1.5 μM PPDA, followed by puffs after bath application of 100 μM D-AP5, a broad NMDAR antagonist. All drugs were washed in for 10 minutes before testing how the drugs affected the response to glutamate puffs.

### Intracranial virus injections

Mice used for intracranial injections of recombinant adeno-associated viruses (AAVs) were between ages P26 – 39. Mice were anesthetized with 1-3% isoflurane (Piramal Critical Care, # NDC 66794-017-25) and body temperature was maintained using a homeothermic heating pad. To minimize postoperative pain, an injection of carprofen (5 mg/kg, CarproJect, Henry Schein Animal Health) was administered subcutaneously. The scalp was shaved using scissors and an incision was made along the midline of the scalp to expose the skull. The injection site was identified using previously validated coordinates for the IC (all coordinates are relative to the skull surface at lambda; in μm: 900 caudal, 1000-1250 lateral, 1850 deep, with two penetrations, one for each lateral coordinate) and a craniotomy was drilled using a micromotor drill (K.1050, Foredom Electric Co.) with a 0.5 mm burr (Fine Science Tools). Glass injection pipettes were pulled from capillary glass (Drummond Scientific Company) using a P-1000 microelectrode puller (Sutter Instruments) and cut on a diagonal for a beveled opening of ∼20 μm. Pipettes were first backloaded with mineral oil and then front filled with AAV1.Syn.Chronos-GFP.WPRE.bGH (Addgene, #59170-AAV1, titer: 1.4e13, 2.2e13), AAV5.Syn.Chronos-GFP.WPRE.bGH (Addgene, #59170-AAV5, titer: 5.3e12), or AAV9.Syn.Chronos-GFP-WPRE.bGH (University of North Carolina Vector Core, Addgene plasmid #59170, titer: 4.5e12) (Klapoetke et al., 2014). The virus was injected using a NanoJect III nanoliter injector (Drummond Scientific Company) connected to an MP-285 micromanipulator (Sutter Instrument). IC injections were made in two penetrations along the medial-lateral axis that were 250 µm apart, and viral deposits were made at 250 µm intervals along the ventral-dorsal axis for a total of 4 deposits at the more medial site and 3 deposits at the more lateral site. 20 nL of virus was deposited per injection, for a total of 150 nL virus injected per IC. After the injections were completed, the scalp was closed either by suturing with Ethilon 6-0 (0.7 metric) nylon sutures (Ethicon USA, LLC) or applying Vetbond (3M, #1469SB) on top of the closed incision. For postoperative analgesia, 0.5 mL of 2% Lidocaine hydrochloride jelly (Akorn Inc) was placed on top of the sutures. Mice were observed for 1 hr for indications of pain or distress and then returned to the vivarium once they were ambulatory. Mice were monitored daily until sutures fell out and the wound was completely healed. Sutures remaining on the 10^th^ post-operative day were manually removed.

### Current-clamp recordings

Slices were placed in a recording chamber and continuously perfused at a rate of ∼2 ml/min with 34 °C oxygenated ACSF. Whole-cell current-clamp recordings were performed with a BVC-700A patch clamp amplifier (Dagan Corporation). Data were low-pass filtered at 10 kHz, sampled at 50 kHz with a National Instruments PCIe-6343 data acquisition board, and acquired using custom software written in Igor Pro. Recording pipettes were pulled from borosilicate glass pipettes (outer diameter 1.5 mm, inner diameter 0.86 mm, Sutter Instrument) with a P-1000 microelectrode puller (Sutter Instrument) and filled with an internal solution containing (in mM): 115 K-gluconate, 7.73 KCl, 0.5 EGTA, 10 HEPES, 10 Na_2_phosphocreatine, 4 MgATP, 0.3 NaGTP, supplemented with 0.1% biocytin (w/v), pH adjusted to 7.3 with KOH and osmolality to 290 mmol/kg with sucrose. All membrane potentials for current-clamp recordings were corrected for an 11 mV liquid junction potential.

### Optogenetics

Current-clamp recordings were conducted 2 – 4 weeks after virus injections to allow time for Chronos expression. Recordings were conducted as described above, except that brain slices were prepared and incubated under red light to limit Chronos activation. To verify the virus injection location in IC injections, fluorescence was visualized under the microscope during the recording session. Recordings were targeted to neurons contralateral to the injection site.

Chronos was activated using 2-10 ms pulses of 470 nm light emitted by a blue LED coupled to the epi-fluorescence port of the microscope and delivered to the slice through a 0.80 NA 40x water immersion objective. Blue light flashes illuminated the entire field of the 0.80 NA 40x objective, corresponding to optical power densities of 6 to 48 mW/mm^2^. Optical power was set using the minimum stimulation needed to elicit an EPSP from the recorded neuron. Recording sweeps with light flashes were repeated 10-30 times with 10 ms between light flashes.

### RNAScope in situ hybridization

Fluorescent in situ hybridization was performed using the RNAscope Multiplex Fluorescent V2 Assay (Advanced Cell Diagnostics, catalog # 320850). Our methods were identical to those previously described (Beebe et al., 2022) and followed manufacturer recommendations (Wang et al., 2012). Briefly, one male (P47) and two female (P45) mice were deeply anesthetized with isoflurane and brains were rapidly removed and frozen on dry ice. Brains were placed in a -80 °C freezer until the day of slicing. Prior to slicing, brains were equilibrated at -20 °C for 1 hour. Brains were sliced into 15 µM sections using a cryostat at -20 °C and sections were mounted on Superfrost Plus slides (Fisher Scientific, catalog # 22037246). For each mouse, five representative sections were chosen spanning the caudal—rostral axis. Sections were fixed using 10% neutral buffered formalin (Sigma-Aldrich, catalog # HT501128) and dehydrated using repeated washes in 50%-100% ethanol. Slides were dried using a Kim wipe and a hydrophobic barrier was drawn around each section. Slices were next incubated in hydrogen peroxide for 10 minutes at room temperature, followed by application of Protease IV for 30 minutes. Probes for *tdTomato*, *Glun2c*, and *Glun2d* (all experimental slices except controls; Advanced Cell Diagnostics catalog #: 317041-C3, 445581-C2, 425951, respectively), along with positive and negative controls (one control slice each), were applied to slices and incubated for 2 hours at 40 °C. The probes were amplified three times for 30 minutes each time at 40 °C using AMP 1-3. The signal was developed using the HRP for each channel and then opal dyes diluted at 1:1000 were assigned for each channel: *tdTomato* was assigned to Opal 690 (Akoya Bioscience, catalog # FP1497001KT), *Glun2c* was assigned to Opal 570 (Akoya Bioscience, catalog # FP1488001KT), and *Glun2d* was assigned to Opal 520 (Akoya Bioscience, catalog # FP1487001KT). Slices were counterstained with DAPI and coverslipped with ProLong Gold antifade mountant (Fisher Scientific, catalog # P36934). Slices were imaged within 1 week of performing the assay using the 0.75 NA 20X objective on a Leica TCS SP8 laser scanning confocal microscope (Leica Microsystems) at 2 µm depth intervals. Emission wavelengths were adjusted for each channel as follows: DAPI (405 nm laser, 410-441 nm), *Glun2d* (488 nm laser, 491-528 nm), *Glun2c* (552 nm laser, 565-583 nm), *tdTomato* (638 nm laser, 690-727 nm).

Co-localization of *Glun2c* and *Glun2d* in *tdTomato*-positive neurons was quantified manually using Neurolucida 360 (MBF Bioscience). One side of the IC (left or right) was selected randomly for quantification in each slice. *tdTomato*-positive neurons were first identified by placing a marker on top of the cell, and then *Glun2c* and *Glun2d* fluorescence was quantified separately for each marked cell. Cells were considered positive for *Glun2c* or *Glun2d* when one or more puncta co-localized with the *tdTomato* fluorescence. All *tdTomato*-positive neurons counted co-labeled with DAPI.

Subdivisions of the IC were determined for each IC slice used in the above analysis by comparison to a reference series of sections from a control C57BL/6J mouse aged P49 that were immunolabeled for GAD67 and GlyT2. This pattern of labeling is routinely used to identify IC subdivisions (Choy Buentello et al., 2015; Beebe and Schofield, 2021), including in our previous studies (Silveira et al., 2020; Anair et al., 2022).

### Analysis of electrophysiological recordings

Amplitude, halfwidth, rise time, and decay time constant measurements were made using custom algorithms in Igor Pro 8. Voltage-clamp data were low-pass filtered at 3 kHz prior to analysis, except for the commissural optogenetic experiment (**Figure 5**), where data were low-pass filtered at 1 kHz and the following function was fit to each response, which was then analyzed:

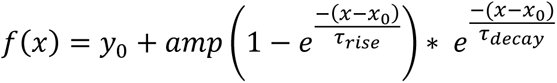

For the temporal summation experiment (**Figure 6**), individual sweeps from each condition were averaged and measurements were made from the average waveform.

### Headbar implantation

Mice used for in vivo experiments were either CBA/CaJ (N = 1 female, age at recording = P51), CBA/C57 (N = 7; 4 females, age range at recording = P81-127), or VIPxAi14 (N = 2 females, ages 50, 53 at recording). Mice were anesthetized with 1-3% isoflurane (Piramal Critical Care, # NDC 66794-017-25) and body temperature was maintained using a homeothermic heating pad. To minimize postoperative pain, injections of carprofen (5 mg/kg, CarproJect, Henry Schein Animal Health) and buprenorphine (0.1 mg/kg) were administered subcutaneously. The scalp was shaved using scissors and an incision was made along the midline of the scalp to expose the skull. The inferior colliculus was identified using previously validated coordinates for either the IC (all coordinates are relative to lambda; in μm: 900 caudal, 1125 lateral) and marked using a surgical marker. The skull was numbed with 0.2 mL bupivacaine and then the periosteum was removed using a scalpel and the skull was scored. After the skull was prepped, a thin layer of Metabond (C&B Metabond from Parkell, cat #s S398, S399, S371) was applied using a fine paintbrush and allowed to dry for 10 minutes. A thin layer of dental acrylic (Henry Schein, Inc., # 1259208 and # 1250143) was then applied on top of the Metabond, and then dental acrylic was applied to a custom-designed headbar which was placed on the skull. Layers of dental acrylic were added to the headbar to secure it to the skull, and then the opening above the IC was filled with silicon elastomer (Smooth-On Inc. Body Double FAST) and secured with a thin strip of dental acrylic. Mice were observed for 1 hr for indications of pain or distress and then returned to the vivarium once they were ambulatory.

For postoperative analgesia, an additional injection of buprenorphine (0.1 mg/kg) was given the day following the surgery, and mice were monitored daily for 7 days.

### In vivo electrophysiological recordings with drug delivery

Mice were allowed to recover for 3 days following headbar implantation and then began habituation to the head fixation setup, which consisted of 1 day of handling within the cage and 3-5 days of head fixation in the sound booth for increasing time duration (10 minutes, 20 minutes, 40 minutes, 60 minutes, 80 minutes). After the habituation period, a craniotomy was surgically drilled above the left IC using the procedures outlined here under “intracranial virus injections”. Mice were allowed to recover for 1 hour, and then placed in the head fixation setup. For drug delivery, drug delivery pipettes were pulled identically to those used for virus injection in the “intracranial virus injections” section. Drug delivery pipettes were affixed at a ∼30 degree angle to recording pipettes (pulled identically to as described in “current clamp recordings”) using melted beeswax (Fisher Scientific, # S25192A) such that the tips of both pipettes were 50-200 µm apart. The drug delivery pipette was backfilled with 20 µM DQP (Tocris, #4491) diluted in the in vivo internal solution (in mM, 1.5 CaCl_2_, 135 NaCl, 1 MgCl_2_, 5.4 KCl, 5 HEPES) and connected to FEP tubing which was attached to a 10 µL Hamilton syringe in a syringe pump (World Precision Instruments Micro4 MicroSyringe Pump Controller). The recording pipette was backfilled with in vivo internal solution and advanced into the brain slowly using an MP-285 micromanipulator (Sutter Instruments). Cells were identified by an increase in resistance and often the presence of spikes in the test pulse. Loose seals (11-30 MΩ) were made to cells using application of a small amount of negative pressure.

Sounds were presented to awake mice using a calibrated, free-field electrostatic speaker (Tucker-Davis Technologies, ES1) positioned ∼10 cm in front of the right ear (contralateral to the IC recording site) at a ∼45 ° angle relative to the midline of the mouse. Sounds consisted of 1 s white noise bursts (4-64 kHz, 70 dB SPL) that were amplitude modulated at 100% modulation depths using modulation frequencies ranging from 16 – 512 Hz at 4 steps/octave. Each stimulus was repeated 8 times in a pseudorandomized order. After collection of the control data, DQP or vehicle solution was infused into the brain at a rate of 0.1 µL/min while the sound presentation was repeated. Mice were recorded from no more than once per day and for fewer than 7 sessions. After recording sessions were finished, the location of the recordings was verified post hoc by dissection.

### Model neuron

The model neuron used here was based, with minimal changes, on the fast-spiking interneuron model described previously by Wang and Buzsáki (Wang and Buzsáki, 1996). Simulated currents interacted with passive membrane properties according to the equation 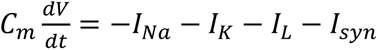 where *C_m_* = 1 µF/cm^2^. Leak current was varied in the model to approximate a range of membrane resistances that might be found in the IC, and the leak reversal potential, *E_L_*, was set to -65 mV.

Action potentials were modeled using Hodgkin-Huxley-type, voltage-dependent Na^+^ and K^+^ conductances (Hodgkin and Huxley, 1952). The transient sodium current is described as 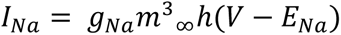 where the activation variable *m* is assumed to be fast, and is thus substituted by its steady state function 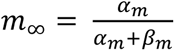 where 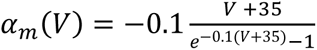 and 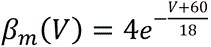. The inactivation variable *h* obeys first order kinetics, where 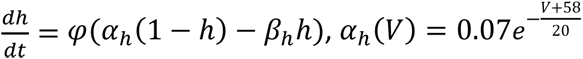, and 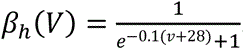. The sodium conductance was set to *g_Na_* = 35 mS/cm^2^ with a reversal potential *E_Na_* = 55 mV.

The delayed rectifier potassium channel was modeled as *I*_*K*_ = *g*_*K*_*n*^4^(*V* − *E*_*K*_), where the activation variable *n* is given by 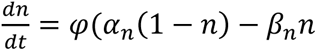. Here, 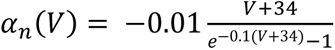 and 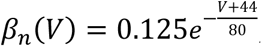, and we set the potassium conductance to *g_K_* = 9 mS/cm^2^ with a reversal potential *E_K_* = -90 mV.

Simulations using the model neuron were performed in MATLAB Version 2022a-2024a (MathWorks) using custom MATLAB scripts adapted from functions provided in (Börgers, Christoph, 2017). All code used for the model simulations can be found at: https://github.com/audreydrotos/NMDA_modeling.

### Statistical analyses

Data analysis was performed using custom algorithms and statistical packages in Igor Pro 8, MATLAB R2021a-2024a (MathWorks), and R 4.1.0 (The R Project for Statistical Computing). Statistical hypothesis testing was performed using two approaches. Comparisons between control and drug condition data for in vitro experiments were made using linear mixed model (LMM) analyses, which were conducted in R using the “lme4” and “lmerTest” packages (Bates et al., 2015; Kuznetsova et al., 2017). Note that in cases where two groups were being compared, the *p*-values from the LMM are identical to those obtained from a paired-samples *t*-test (Shravan Vasishth et al., 2022). For comparisons between control and vehicle or drug conditions in vivo, we used a Wilcoxon Signed Rank test, which was performed using the ‘signrank’ function in MATLAB. Effects were considered significant when *p* < 0.05.

For in vitro electrophysiology data involving repeated measures from individual cells, effect sizes and confidence intervals are shown using Gardner-Altman estimation plots (two groups) or Cumming estimation plots (three groups). These plots were made using custom MATLAB scripts modeled after the DABEST package (Ho et al., 2019). Both types of estimation plots use parallel coordinate plots to show the responses of individual cells for each condition, and these are accompanied by histograms showing the range and 95% confidence intervals of the paired mean differences expected between each drug condition and the control based on a bootstrap analysis. Our approach to generating these plots has been detailed previously (Rivera-Perez et al., 2021) and was influenced by the estimation statistics approach (Bernard, 2019; Calin-Jageman and Cumming, 2019). While estimation plots are provided to give additional insight into the data, all determinations of statistical significance were based on the standard null-hypothesis statistical tests detailed in the preceding paragraph.

## Results

### NMDAR currents in VIP neurons are sensitive to GluN2C/D-specific pharmacology

Several previous studies have noted that NMDARs often contribute to subthreshold postsynaptic responses in IC neurons (Kelly and Zhang, 2002; Ma et al., 2002; Wu, 2004; Oberle et al., 2022) . Similarly, we recently found that NMDARs contributed to subthreshold EPSPs elicited by commissural inputs to IC VIP neurons (Goyer et al., 2019). NMDAR currents at resting potential are unusual, as most NMDARs in the adult brain have a voltage-dependent Mg^2+^ block which prevents ions from flowing through the channel pore at or near resting potential (Traynelis et al., 2010; Paoletti et al., 2013). However, sensitivity to Mg^2+^ block depends on the subunit composition of the receptor. While receptors containing GluN2A/B subunits are highly sensitive to Mg^2+^ block, receptors containing GluN2C/D subunits are less sensitive, allowing them to conduct current even at hyperpolarized potentials (Siegler Retchless et al., 2012). Based on this, we hypothesized that VIP neurons express NMDARs containing GluN2C and/or GluN2D subunits.

To test this hypothesis, we voltage-clamped VIP neurons at -70 mV and used a puffer pipette containing 1 mM glutamate to elicit EPSCs (**Figure 1A**). We found that 10 ms glutamate puffs reliably elicited EPSCs in VIP neurons (**Figure 1B**). We next performed puffs in the presence of 1.5 µM PPDA, a GluN2C/D-selective antagonist (Lozovaya et al., 2004; Li et al., 2020; Jing et al., 2022), and then in the presence of 20 µM CIQ, a GluN2C/D-selective positive allosteric modulator (Mullasseril et al., 2010; Feng et al., 2014; Zhang et al., 2014; Nouhi et al., 2018; Liu et al., 2021). PPDA significantly decreased the amplitude (LMM: *p* = 0.038) of the elicited EPSC. Our experimental design did not favor large CIQ effects since we used a saturating glutamate concentration (1 mM) in our puff solution. At the population level, CIQ did not alter the amplitude (*p* = 0.088), halfwidth (*p* = 0.15), rise time (*p* = 0.49), or decay tau (*p* = 0.10) of the EPSC (**Figure 1C**). The sensitivity of NMDAR currents in VIP neurons to PPDA and, in some cases to CIQ, suggests that at least a portion of the NMDARs expressed in VIP neurons contain GluN2C/D subunits.

**Figure 1.**
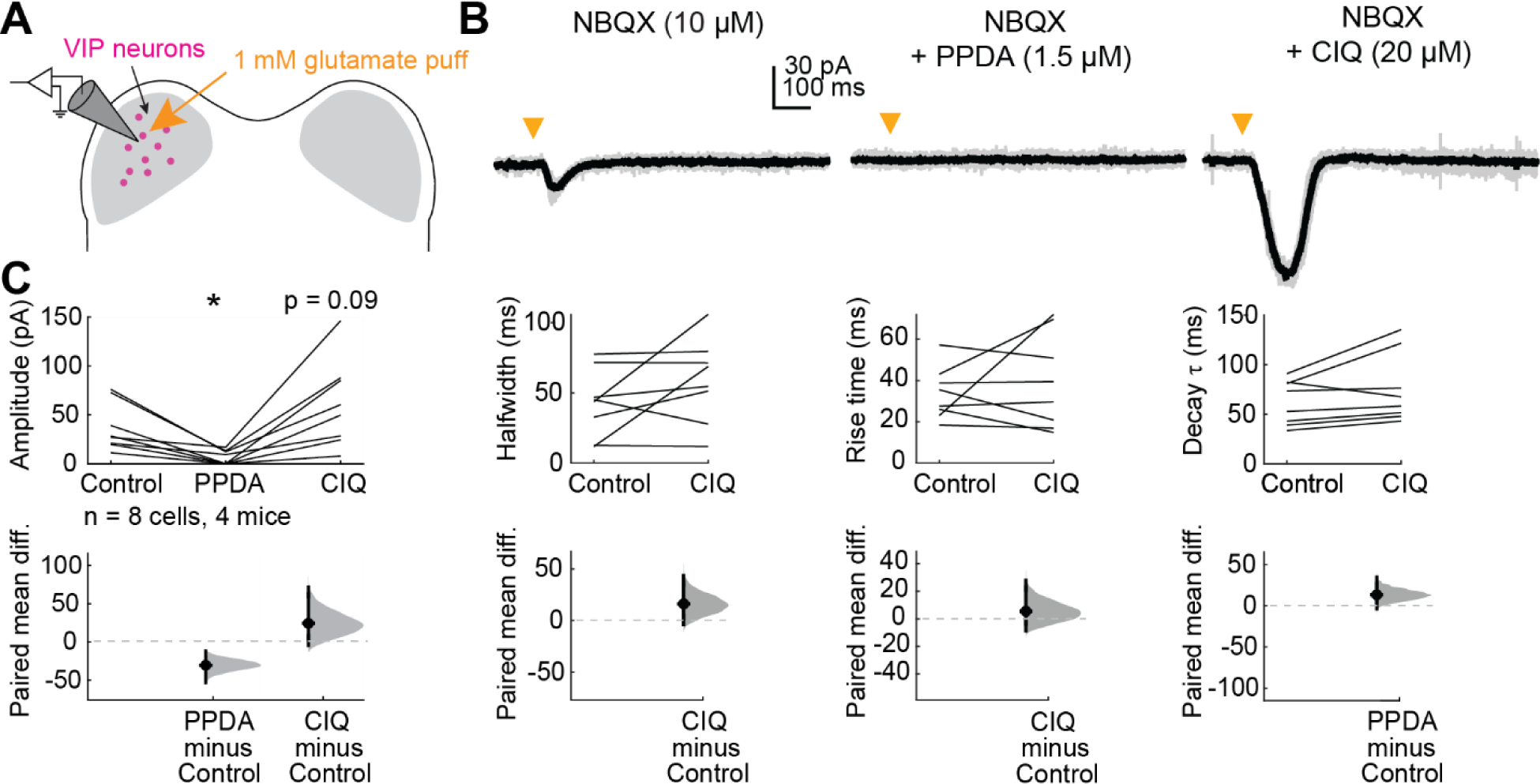
VIP neurons express GluN2C/D-containing NMDARs. ***A,*** Puffs of glutamate onto VIP neurons activated NMDARs. Pharmacology was used to examine whether GluN2C/D-containing receptors contributed to elicited currents. ***B,*** Brief (10 ms) puffs of glutamate onto a VIP neuron elicited a small inward current (left) which was completely abolished by the GluN2C/D antagonist PPDA (middle) and enhanced by the GluN2C/D positive allosteric modulator CIQ (right). ***C,*** PPDA decreased the amplitude (LMM: β = -30.46, 95% CI [-56.32, -4.61], *p* = 0.038, *n* = 8) of the response to the glutamate puff. CIQ trended towards enhancing the amplitude (β = 24.33, 95% CI [-1.53, 50.18], *p* = 0.088, *n* = 8) of the response, but did not alter the halfwidth (β = 16.13, 95% CI [-7.89, 37.16], *p* = 0.15, *n* = 8), rise time (β = 5.56, 95% CI [-10.24, 21.36], *p* = 0.49, *n* = 8) or decay τ (β = 13.23, 95% CI [-1.29, 27.74], *p* = 0.10, *n* = 8) of the response to the glutamate puff. Vertical black lines on the paired mean difference plots indicate 95% bootstrap confidence intervals, and gray histograms show the expected distribution of paired mean differences based on bootstrap analysis.

### Many IC neurons in adult mice express *Glun2c* and/or *Glun2d* mRNA, with VIP neurons predominately expressing *Glun2d*

GluN2C and GluN2D subunits are similar in their sensitivity to Mg^2+^ block, but expression levels of *Glun2c* and *Glun2d* mRNA change dramatically and in different directions during development (Paoletti et al., 2013). GluN2D-containing NMDARs (along with GluN2B-containing NMDARs) are primarily found early in development (Wenzel et al., 1996) and are largely replaced with GluN2A and GluN2C subunits by adulthood (Paoletti et al., 2013), with some exceptions in small populations of interneurons, such as those found in hippocampus (Monyer et al., 1992). In many cases, the developmental switch in NMDAR subunit composition is thought to be driven by sensory experience (Traynelis et al., 2010). Expression of GluN2 subunits is also spatially regulated: while GluN2A and GluN2B-containing NMDARs are typically found at the synapse, GluN2C and GluN2D-containing NMDARs are often found at extrasynaptic sites (Paoletti et al., 2013). However, growing evidence points to GluN2C/D expression at synapses, such as in striatum (Logan et al., 2007) and substantia nigra (Brothwell et al., 2008).

Based on these precedents, we hypothesized that IC VIP neurons in adult animals express GluN2C subunits. To test this hypothesis, we used the RNAScope assay to perform fluorescence in situ hybridization on IC brain slices from three VIP-IRES-Cre x Ai14 mice aged P45-47 (**Figure 2A,B**). Using probes for *tdTomato*, *Glun2c*, and *Glun2d* mRNA, we were surprised to find that 91.4% of VIP neurons expressed *Glun2d* and only 8.1% expressed *Glun2c*. More specifically, 84.3% of VIP neurons expressed only *Glun2d*, 1.0% of VIP neurons expressed only *Glun2c*, and 7.1% of VIP neurons expressed both. The remaining 7.7% of VIP neurons did not express *Glun2c* or *Glun2d* (**Table 1**, **Figure 3A**).

**Figure 2.**
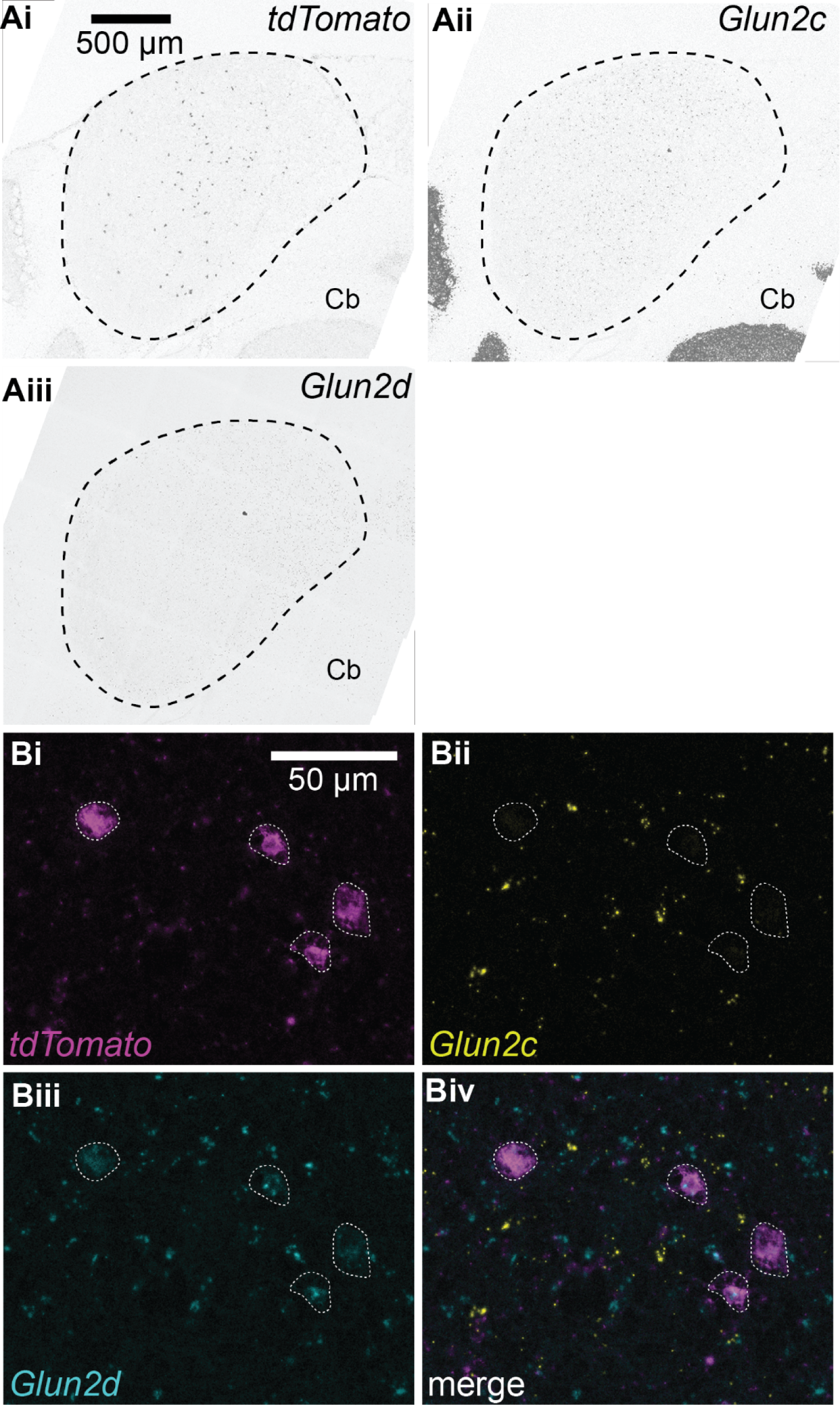
*Glun2c and Glun2d* mRNA is widely expressed throughout the IC, and most VIP neurons express *Glun2d*. ***A,*** 40x confocal images of IC brain slices following RNAscope in situ hybridization showing expression of *tdTomato* mRNA ***(i)***, which is expressed by VIP neurons in VIP-IRES-Cre x Ai14 mice, *Glun2c* mRNA ***(ii)***, and *Glun2d* mRNA ***(iii)***. ‘Cb’ indicates portions of the cerebellum that were captured in the images. ***B,*** 63x confocal images with white arrows indicating *tdTomato*-positive neurons ***(i)***. 8.1% of *tdTomato* positive neurons expressed *Glun2c* mRNA ***(ii)*** and 91.4% expressed *Glun2d* mRNA ***(iii)***. *Glun2c* and *Glun2d* mRNA was also commonly observed in tdTomato-negative neurons, suggesting that these NMDAR subunits are expressed in multiple IC neuron types. A merge of these images is shown in ***iv***.

**Figure 3.**
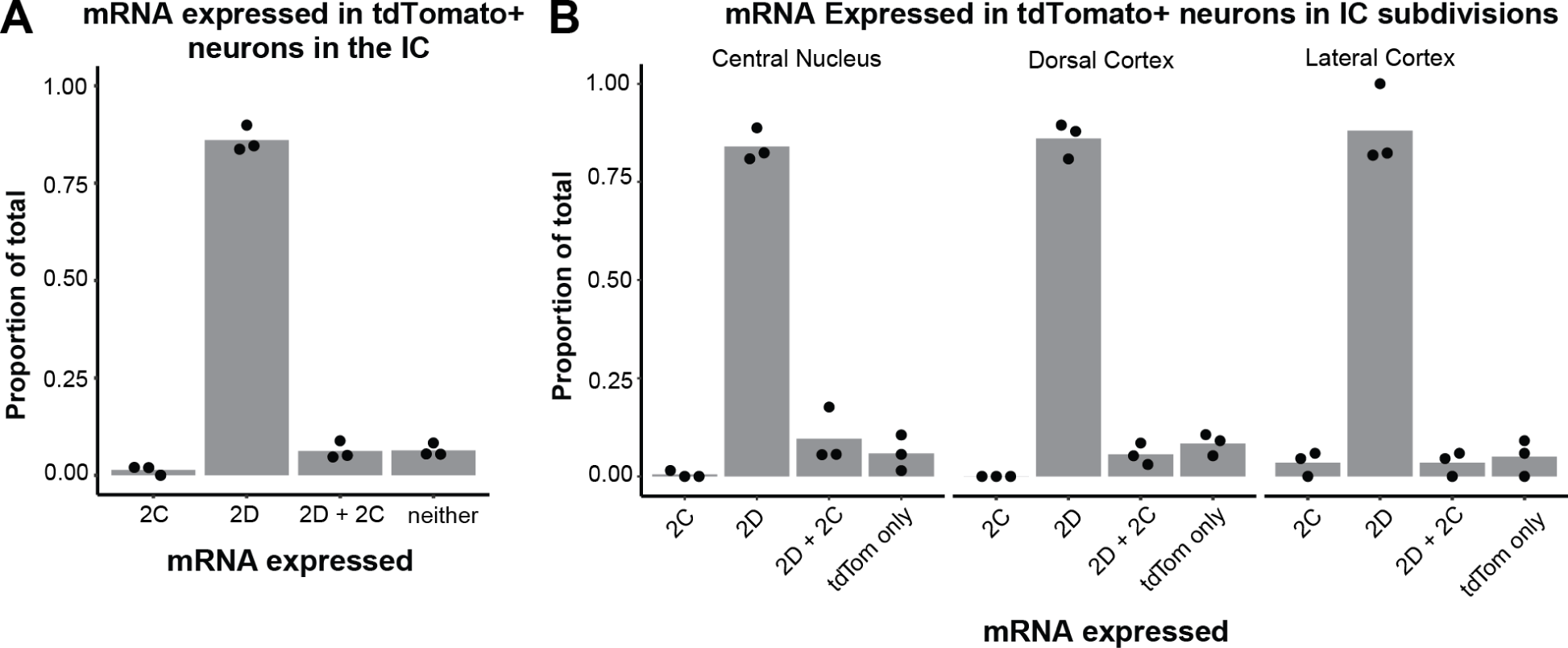
91.4% of VIP neurons express *Glun2d* mRNA. ***A,*** Quantification of mRNA expressed in *tdTomato*-positive (VIP) neurons in the IC. 1.0% of VIP neurons expressed only *Glun2c* mRNA, while 84.3% of VIP neurons expressed only *Glun2d* mRNA. 7.1% of VIP neurons expressed both *Glun2c* and *Glun2d* mRNA. 7.7% of VIP neurons did not express *Glun2c* or *Glun2d* mRNA. Individual data points represent the mean for individual mice. ***B,*** The central nucleus, lateral cortex, and dorsal cortex of the IC did not differ in their expression profiles of tdTomato-positive neurons expressing only *Glun2c* (0.8%, 0.0%, 3.8%, respectively), only *Glun2d* (83.7%, 85.0%, 86.5%), both *Glun2c* and *Glun2d* (7.9%, 6.2%, 3.8%), and neither *Glun2c*/*Glun2d* (7.6%, 8.8%, 5.8%). Individual data points represent the means for individual mice.

**Table 1.** Percentage of *tdTomato+* neurons expressing *Glun2d* and/or *Glun2c* mRNA. The number and percentage of *tdTomato* positive cells that expressed *Glun2c* and *Glun2d* mRNA using an RNAScope in situ hybridization assay. Data is from a series of coronal IC sections collected from three VIP-IRES-Cre x Ai14 mice aged P45-47.

| Animal | Slice (n) | No. of *tdT*<sup>+</sup> | No. of *Gln2d*<sup>+</sup> | No. of *Gln2c*<sup>+</sup> | No. of *tdT*<sup>+</sup> *Gln2d*<sup>+</sup> *Gln2c*<sup>+</sup> | % *tdT*<sup>+</sup> *Gln2d*<sup>+</sup> | % *tdT*<sup>+</sup> *Gln2c*<sup>+</sup> | % *tdT*<sup>+</sup> *Gln2d*<sup>+</sup> *Gln2c*<sup>+</sup> |
| --- | --- | --- | --- | --- | --- | --- | --- | --- |
| **Mouse 1** | A caudal | 42 | 36 | 3 | 3 | 85.7% | 7.1% | 7.1% |
|  | B caudal | 52 | 49 | 4 | 4 | 94.2% | 7.7% | 7.7% |
|  | C caudal | 48 | 47 | 2 | 2 | 97.9% | 4.2% | 4.2% |
|  | D rostral | 3 | 3 | 0 | 0 | 100.0% | 0.0% | 0.0% |
|  | E rostral | 4 | 4 | 0 | 0 | 100.0% | 0.0% | 0.0% |
|  | **Total** | **149** | **139** | **9** | **9** | **93.3%** | **6.0%** | **6.0%** |
| **Mouse 2** | D caudal | 71 | 68 | 9 | 9 | 95.8% | 12.7% | 12.7% |
|  | B medial | 13 | 10 | 2 | 1 | 76.9% | 15.4% | 7.7% |
|  | C medial | 26 | 26 | 4 | 4 | 100.0% | 15.4% | 15.4% |
|  | A rostral | 3 | 3 | 0 | 0 | 100.0% | 0.0% | 0.0% |
|  | **Total** | **113** | **107** | **15** | **14** | **94.7%** | **13.3%** | **12.4%** |
| **Mouse 3** | B caudal | 94 | 83 | 10 | 7 | 88.3% | 10.6% | 7.4% |
|  | D caudal | 85 | 76 | 5 | 5 | 89.4% | 5.9% | 5.9% |
|  | A medial | 70 | 62 | 2 | 1 | 88.6% | 2.9% | 1.4% |
|  | C rostral | 10 | 9 | 1 | 1 | 90.0% | 10.0% | 10.0% |
|  | **Total** | **259** | **230** | **18** | **14** | **88.8%** | **6.9%** | **5.4%** |
| **Grand total** |  | **521** | **476** | **42** | **37** | **91.4%** | **8.1%** | **7.1%** |

We also investigated whether NMDAR subunit expression differed among VIP neurons present in the three major IC subdivisions: central nucleus, dorsal cortex, and lateral cortex. We found that the percentage of VIP neurons expressing various NMDAR subunits was similar across all IC subdivisions (**Table 2**, **Figure 3B**). These results suggest that VIP neurons preferentially express GluN2D rather than GluN2C subunits, regardless of their location in the IC.

**Table 2.**
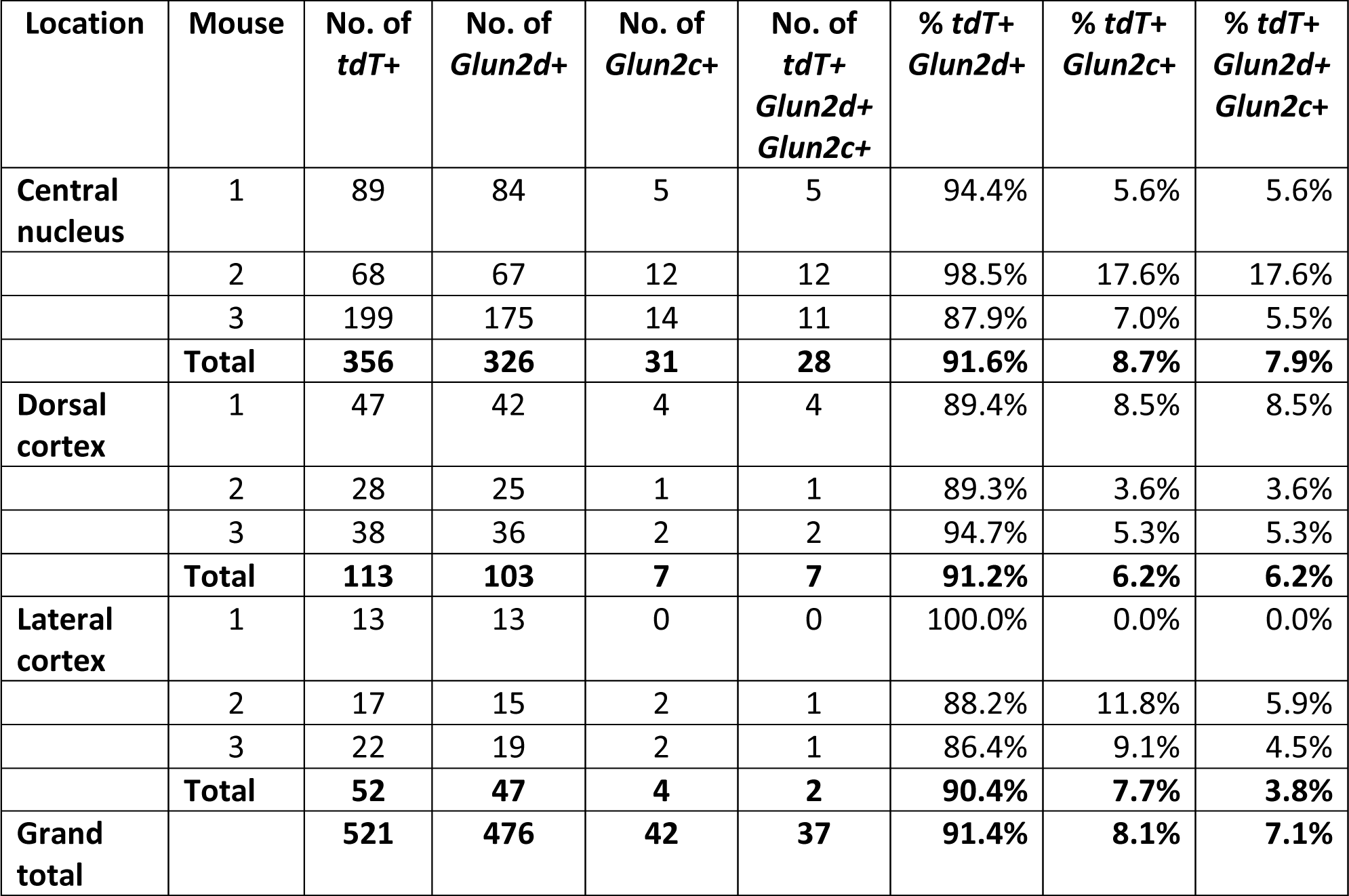
Percentage of neurons that expressed *Glun2d* and *Glun2c* in IC central nucleus, dorsal cortex, and lateral cortex. The number and percentage of *tdTomato* positive cells in the three major IC subdivisions (central nucleus, lateral cortex, and dorsal cortex) that expressed *Glun2c* and *Glun2d* mRNA using an RNAScope in situ hybridization assay. Data is from a series of coronal IC sections collected from three VIP-IRES-Cre x Ai14 mice aged P45-47.

| Location | Mouse | No. of *tdT+* | No. of *Glnu2d+* | No. of *Glnu2c+* | No. of *tdT+ Glnu2d+ Glnu2c+* | % *tdT+ Glnu2d+* | % *tdT+ Glnu2c+* | % *tdT+ Glnu2d+ Glnu2c+* |
| --- | --- | --- | --- | --- | --- | --- | --- | --- |
| Central nucleus | 1 | 89 | 84 | 5 | 5 | 94.4% | 5.6% | 5.6% |
|  | 2 | 68 | 67 | 12 | 12 | 98.5% | 17.6% | 17.6% |
|  | 3 | 199 | 175 | 14 | 11 | 87.9% | 7.0% | 5.5% |
|  | **Total** | **356** | **326** | **31** | **28** | **91.6%** | **8.7%** | **7.9%** |
| Dorsal cortex | 1 | 47 | 42 | 4 | 4 | 89.4% | 8.5% | 8.5% |
|  | 2 | 28 | 25 | 1 | 1 | 89.3% | 3.6% | 3.6% |
|  | 3 | 38 | 36 | 2 | 2 | 94.7% | 5.3% | 5.3% |
|  | **Total** | **113** | **103** | **7** | **7** | **91.2%** | **6.2%** | **6.2%** |
| Lateral cortex | 1 | 13 | 13 | 0 | 0 | 100.0% | 0.0% | 0.0% |
|  | 2 | 17 | 15 | 2 | 1 | 88.2% | 11.8% | 5.9% |
|  | 3 | 22 | 19 | 2 | 1 | 86.4% | 9.1% | 4.5% |
|  | **Total** | **52** | **47** | **4** | **2** | **90.4%** | **7.7%** | **3.8%** |
| Grand total |  | **521** | **476** | **42** | **37** | **91.4%** | **8.1%** | **7.1%** |

Importantly, our results also show that *Glun2c* and *Glun2d* mRNA is widely expressed in the IC (**Figure 2A**). This suggests that *Glun2c* and *Glun2d*-containing NMDARs are a prominent feature in the IC and likely have a significant and widespread impact on synaptic computations.

### VIP neurons also express GluN2A/B-containing NMDARs

We next investigated whether IC VIP neurons also express GluN2A or GluN2B-containing receptors. In the adult brain, GluN2A-containing NMDARs are more commonly found at synapses and confer more typical properties of the NMDA receptor, including sensitivity to Mg^2+^ block at rest (Siegler Retchless et al., 2012). To determine whether VIP neurons express GluN2A/B-containing NMDARs in addition to GluN2C/D-containing NMDARs, we targeted VIP neurons for whole-cell recordings and puffed glutamate in the presence of AMPAR antagonist NBQX (10 µM, **Figure 4A**). To examine the contributions of all NMDARs to the resulting current, we removed Mg^2+^ from the ACSF and replaced it with equimolar Ca^2+^ to maintain the overall concentration of divalent ions. We found that puffs of glutamate elicited much larger currents in the Mg^2+^-free condition than we observed in the earlier puff experiments (compare **Figures 1C, 4C**).

**Figure 4.**
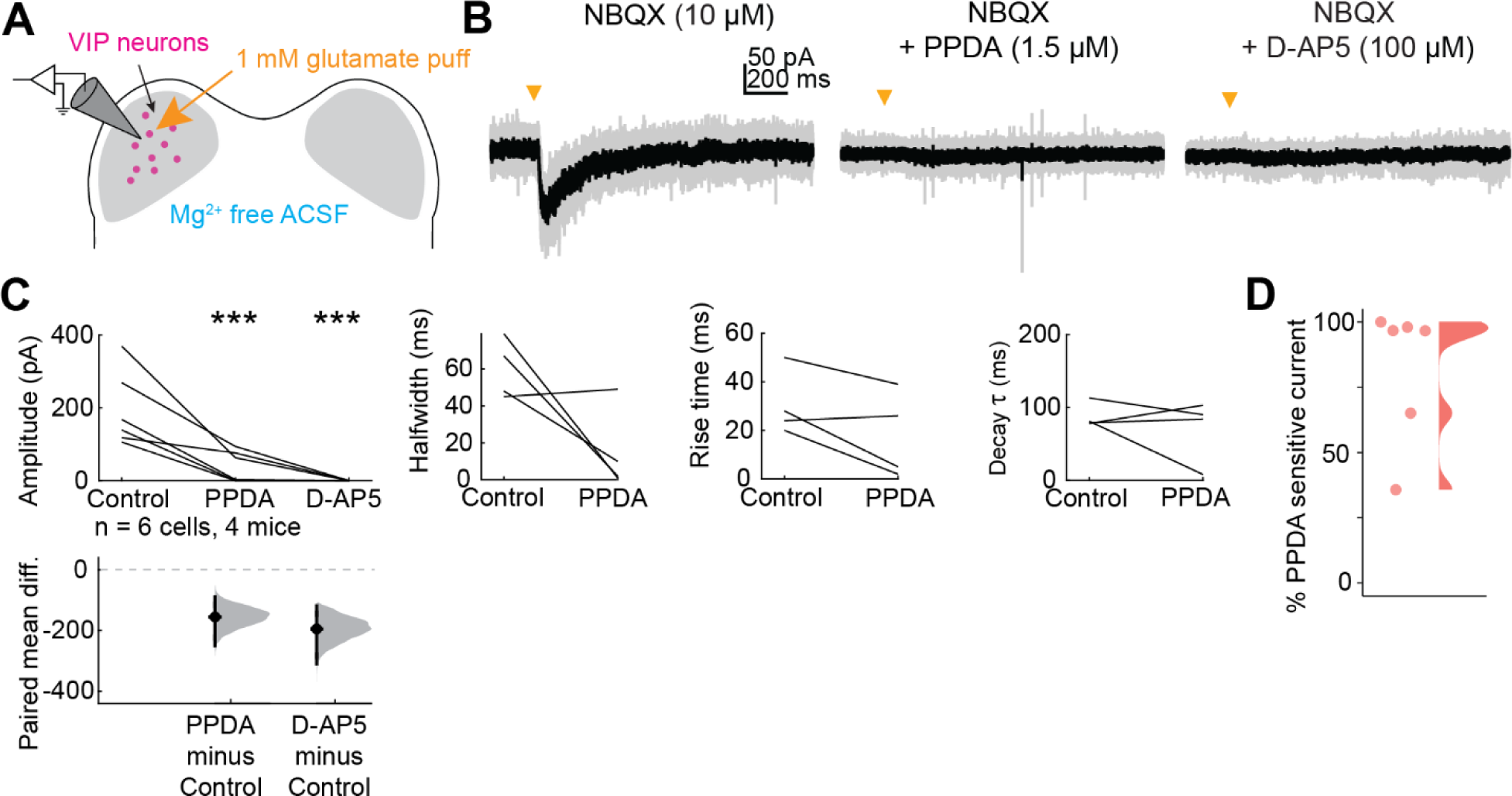
VIP neurons express GluN2A/B-containing NMDARs in addition to GluN2C/D-containing NMDARs. ***A,*** Puffs of glutamate onto VIP neurons activated NMDARs. Pharmacology was used to examine the proportion of GluN2A/B and GluN2C/D containing receptors present on VIP neurons. ***B,*** An EPSC elicited by a brief (10 ms) puff of glutamate onto a VIP neuron was abolished by application of the GluN2C/D antagonist PPDA. ***C,*** PPDA decreased the amplitude (β = -155.49, 95% CI [-220.62, -90.36], *p* = 9.26e-4, *n* = 6) of the response to the glutamate puff. D-AP5 completely abolished the response in all cells tested (amplitude, LMM: β = -194.83, 95% CI [-259.96, -129.70], *p* = 1.71e-4, *n* = 6). Effects on EPSC kinetics for cells with residual current after PPDA application are shown in ***C,*** right panels. Vertical lines on the paired mean difference plots indicate 95% bootstrap confidence intervals, and gray histograms show the expected distribution of paired mean differences based on bootstrap analysis. ***D,*** Percent of EPSC amplitude that was sensitive to PPDA application in each of the 6 cells from **C**. The distribution was created using the ‘ggdist’ and ‘stat_halfeye’ functions from the ‘ggplot’ package for R, which uses a kernel density estimate to generate a curve that reflects the distribution of the data (Kay, 2024).

**Figure 5.**
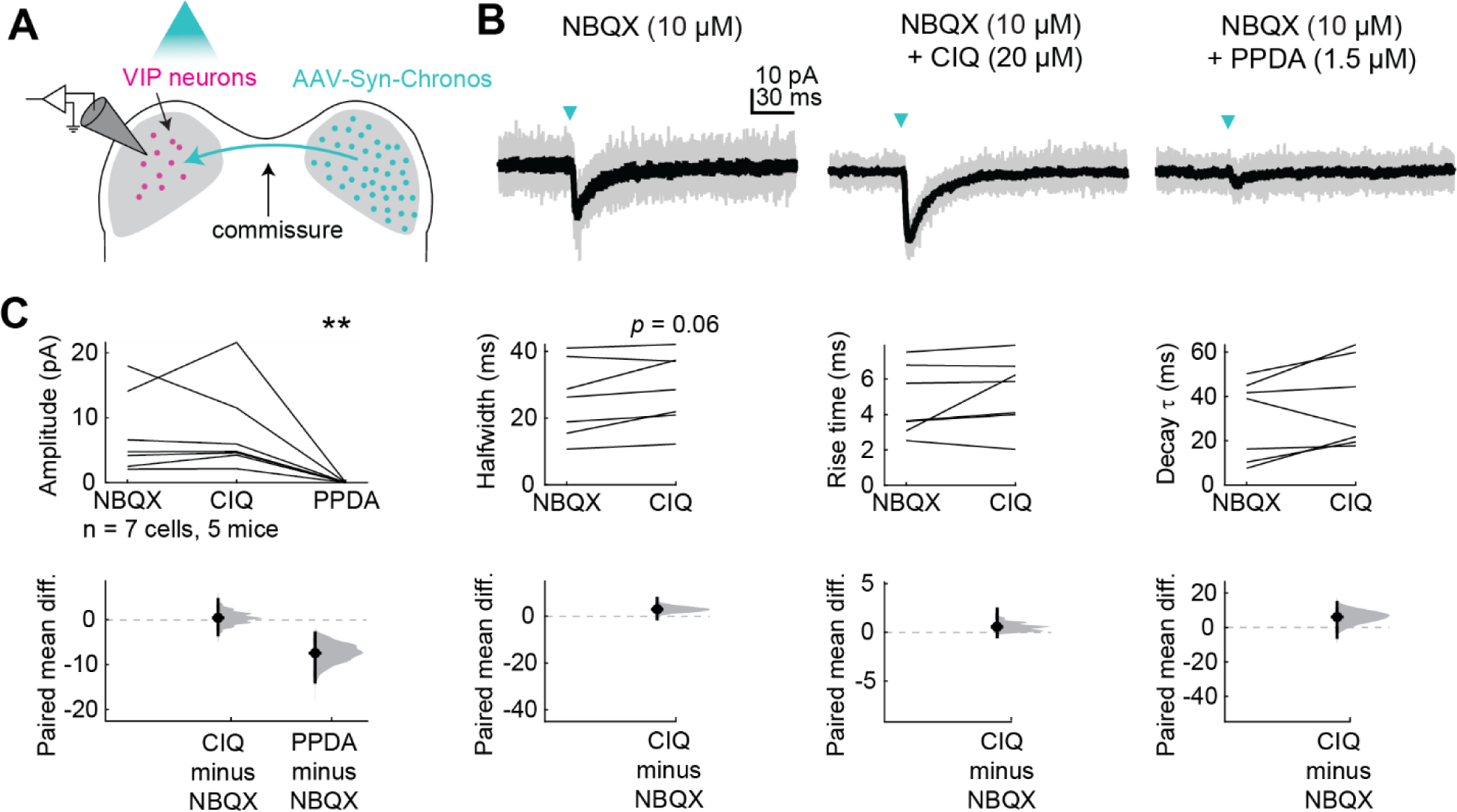
Commissural inputs to VIP neurons activate GluN2C/D-containing NMDARs at resting potential. ***A,*** Viral injections in the right IC drove Chronos expression in commissural projections to the left IC. Optogenetic circuit mapping and pharmacology were used to examine postsynaptic responses in VIP neurons in the left IC. ***B,*** An EPSC elicited by a brief (10 ms) flash of light (left) was enhanced by the GluN2C/D-specific positive allosteric modulator CIQ (middle) and diminished by the GluN2C/D-specific antagonist PPDA (right). ***C,*** CIQ did not significantly alter the amplitude (LMM: β = 0.39, 95% CI [4.63, -3.86], *p* = 0.86, *n* = 7), halfwidth (β = 2.95, 95% CI [0.22, 5.69, *p* = 0.064], *p* = 0.42, *n* = 7), rise time (β = 0.56, 95% CI [-0.38, 1.50], *p* = 0.26, *n* = 7), or decay τ (β = 6.13, 95% CI [-2.02, 14.27], *p* = 0.16, *n* = 7) of EPSCs. PPDA abolished the response in all cells tested (amplitude, LMM: β = -7.44, 95% CI [-11.68, -3.20], *p* = 5.13e-3, *n* = 7). Vertical lines on the paired mean difference plots indicate 95% bootstrap confidence intervals, and gray histograms show the expected distribution of paired mean differences based on bootstrap analysis.

**Figure 6.**
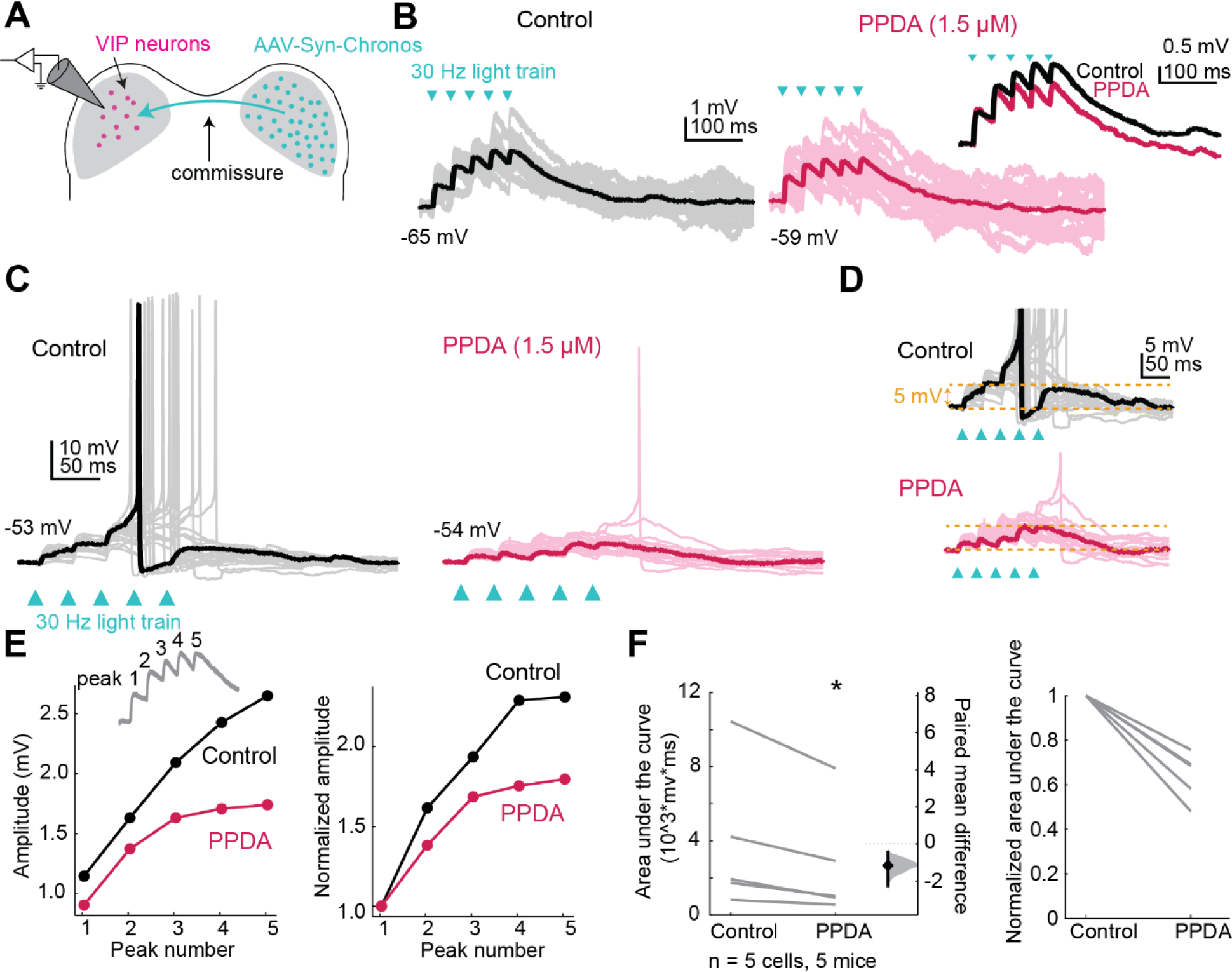
GluN2C/D-containing NMDARs facilitate temporal summation in VIP neurons. ***A,*** Viral injections in the right IC drove Chronos expression in commissural projections to the left IC. Trains of optogenetic stimuli combined with pharmacology were used to examine the contributions of GluN2C/D-containing receptors to temporal summation. ***B,*** EPSP summation elicited by a 30 Hz train of light pulses (left) was decreased by application of the GluN2C/D-specific antagonist PPDA (right, comparison inset). ***C,*** Example of a cell in which summation of optogenetically elicited EPSPs reached action potential threshold, eliciting trains of action potentials. PPDA eliminated spiking in all but one sweep in this cell. ***D,*** Example traces from ***C*** scaled to demonstrate the differences in temporal summation between control and PPDA conditions. ***E,*** The peak amplitude of each EPSP was calculated from the baseline (see inset). The amplitude of the response increased with increasing peak number (LMM: β = 1.06, 95% CI [0.66, 1.47], *p* = 9.15e-6, *n* = 5), although the interaction between peak and drug condition was not statistically significant (β = -0.48, 95% CI [-1.06, 0.095], *p* = 0.11, *n* = 5). ***F,*** PPDA significantly decreased the area under the curve for the train of EPSPs (LMM: β = -116.57, 95% CI [-198.93, - 34.21], *p* = 0.038, *n* = 5), which can be seen in both the non-normalized (left) and normalized (right) plots. Vertical lines on the paired mean difference plots indicate 95% bootstrap confidence intervals, and gray histograms show the expected distribution of paired mean differences based on bootstrap analysis.

Next, we bath applied 1.5 µM PPDA, a GluN2C/D-specific antagonist, and as expected, we found that this significantly decreased the amplitude (LMM: *p* = 9.26e-4), halfwidth (*p* = 8.6e-4), rise time (*p* = 8.45e-3), and decay time constant (*p* = 0.028) of the response, again suggesting that a portion of the NMDAR current in VIP neurons is mediated by GluN2C/D-containing NMDARs (**Figure 4C**). Next, we bath applied 100 µM D-AP5, an NMDAR antagonist that blocks NMDARs regardless of GluN2 subunit identity and found that this completely abolished the response in all neurons (amplitude, *p* = 1.71e-4). Overall, 5 out of 6 VIP neurons had both PPDA sensitive and Insensitive currents, indicating activation of both GluN2A/B and GluN2C/D currents. On average in these cells, 82.0% ± 26.2% of the current was sensitive to PPDA (mean ± standard deviation, **Figure 4D**). Two neurons had current that was completely abolished by PPDA indicating exclusive activation of GluN2C/D-containing receptors. These results suggest that individual VIP neurons can express NMDARs that differ by subunit composition and are thus specialized for different roles in the cell.

### Commissural inputs to VIP neurons activate GluN2C/D-containing NMDARs

We previously found that inputs from the contralateral IC via the IC commissure activate NMDARs on VIP neurons at resting potential (Goyer et al., 2019). To test whether these responses were also mediated by GluN2C/D-containing NMDAR receptors, we repeated these experiments in the presence of GluN2C/D-specific drugs. Commissural projections were labeled with the excitatory opsin Chronos by injecting AAV-Synapsin-Chronos-GFP (serotype 1, 5, or 9) into the right IC of VIP-IRES-Cre x Ai14 mice. 1.5 to 4 weeks later, we targeted whole-cell recordings to fluorescently labeled VIP neurons in the IC contralateral to the injection site (**Figure 5A**). We found that even in the presence of AMPAR antagonist NBQX (10 µM), flashes of blue light elicited small EPSCs (-2 to -18 pA, **Figure 5B**). The GluN2C/D-specific positive allosteric modulator CIQ did not alter the light-evoked response, again possibly because a saturating concentration of glutamate was used and/or because CIQ is less effective at tri-heteromeric NMDARs that contain a GluN2A/B subunit in addition to a GluN2C/D subunit (Mullasseril et al., 2010) (**Figure 5C**, LMM: amplitude *p* = 0.86, halfwidth *p* = 0.064, rise time *p* = 0.26, decay tau *p* = 0.17), but the GluN2C/D-specific antagonist PPDA completely abolished the response in all cells tested (amplitude, LMM: *p* = 5.13e-3). These results support the hypothesis that commissural inputs to VIP neurons activate NMDARs containing GluN2C/D subunits.

### Activation of GluN2C/D-containing NMDARs facilitates temporal summation

Since NMDARs with GluN2C/D subunits have slower kinetics and conduct more current at resting membrane potential than NMDARs with GluN2A/B subunits (Paoletti et al., 2013), expression of GluN2C/D-containing NMDARs could have a particularly strong effect on the duration of excitation elicited by glutamatergic transmission. We therefore hypothesized that GluN2C/D-containing NMDARs in the IC facilitate temporal integration by widening the time window for integration of synaptic inputs. To test this hypothesis, we used optogenetics to stimulate commissural inputs to VIP neurons (**Figure 6A**). Trains of five light pulses at 30 Hz elicited temporal summation in all the cells tested, with the second through fifth EPSPs starting at a more depolarized potential than the first EPSP (**Figure 6B**). To test whether GluN2C/D-containing receptors facilitate this temporal summation, we next elicited trains of input in the presence of the GluN2C/D-containing NMDAR antagonist PPDA (1.5 µM). In one cell, the light train elicited temporal summation great enough to elicit an action potential in the control condition, and this was impaired in the PPDA condition, where the cell only reached action potential threshold in one trial (**Figure 6C,D**). In trials that did not reach action potential threshold, we assessed temporal summation by comparing the peak amplitudes of each of the five EPSPs elicited by the train stimuli in the control and PPDA conditions (**Figure 6B**). The amplitude of the peaks significantly increased during the train (LMM: *p* = 9.15e-6, *n* = 5, **Figure 6E**) indicating temporal summation occurred during the train. This change in amplitude was not correlated with small changes in the resting membrane potential over the course of the recording (r^2^ = 0.001). In addition, PPDA significantly decreased the area under the curve of the train response (LMM: *p* = 0.038, **Figure 6F**), indicating a decrease in temporal summation when GluN2C/D-containing receptors were blocked. These results show that GluN2C/D-containing NMDARs strongly enhance the time window for synaptic integration in VIP neurons.

### GluN2C/D-containing NMDA receptors provide additive gain to rate coding in a neuron model

Our in vitro data suggest that GluN2C/D-containing receptors facilitate synaptic integration in IC neurons, and we hypothesized that this boosts gain in the IC through an increase in temporal summation, which could in turn promote rate coding for auditory stimuli. To test this hypothesis, we created a neuron model based on a previously published Hodgkin-Huxley model of fast-spiking hippocampal interneurons (Wang and Buzsáki, 1996). We based our model on the Wang and Buzsáki model because it provides a well-known and widely used model for neurons that exhibit sustained firing over a large range of synaptic input frequencies, similar to what we observe for VIP neurons in vitro (Goyer et al., 2019). In our adapted model, we included two synaptic conductances: one with kinetics similar to AMPA receptors on VIP neurons, and one with kinetics similar to a GluN1-GluN2D-GluN2D NMDA receptor (Yi et al., 2019). We set the maximal NMDA conductance to 25% of the maximal AMPA conductance and set the leak conductance to 0.7 mS/cm^2^ to mimic the average membrane resistance of a VIP neuron (∼200 MΩ), assuming a 15 µm diameter soma (Goyer et al., 2019). We next simulated trains of synaptic inputs at frequencies ranging from 16 – 512 Hz, activating either the AMPA conductance alone or both the AMPA and NMDA conductances.

We found that activation of the AMPA-only synapse by a slow train of stimuli (16 Hz) elicited a series of phase-locked EPSPs in the model neuron (**Figure 7A**, left panel). When the NMDA conductance was added, the slow kinetics of the NMDA receptor caused the EPSPs to temporally sum across stimulus presentations, providing an additive gain that pushed the neuron over action potential threshold and elicited phase-locked action potentials (**Figure 7A**, right panel). When the input frequency was increased to 64 Hz, the neuron fired phase-locked action potentials in the AMPA-only condition (**Figure 7B**, left panel), and addition of the NMDA conductance again increased the overall firing rate (**Figure 7B**, right panel). To better examine how the NMDA conductance shapes the response to inputs at different stimulus frequencies, we simulated the response using input frequencies from 16 – 512 Hz (4 steps/octave) in both the AMPA-only and AMPA plus NMDA conditions. We found that addition of the NMDA conductance caused an overall increase in firing rate in response to all input frequencies tested (**Figure 7C**, orange line). Addition of the NMDA conductance also altered phase-locking to the response (**Figure 7D**): in the AMPA-only condition, the neuron exhibited near perfect phase locking as soon as the input frequency was high enough to elicit spiking (blue line). When the NMDA conductance was added (orange line), phase locking was high at lower input frequencies but quickly became poorer as the input frequency increased, an effect due in part to the slow kinetics of the NMDA receptor which decreased the ability of the fast, temporally precise AMPA conductance to facilitate phase-locking in the model neuron. In line with this, addition of the NMDA conductance shifted the phase of the input response at which the neuron fired (**Figure 7E**). These results suggest that GluN2C/D-type NMDA receptors provide additive gain to neuron input-output functions, in the form of an upward shift along the *y*-axis of firing vs stimulus curves (e.g., **Figure 7C**) (Silver, 2010). Our model also indicates that, as stimulus frequencies increase, this type of additive gain increases firing rate while simultaneously decreasing the precision of spike timing.

**Figure 7.**
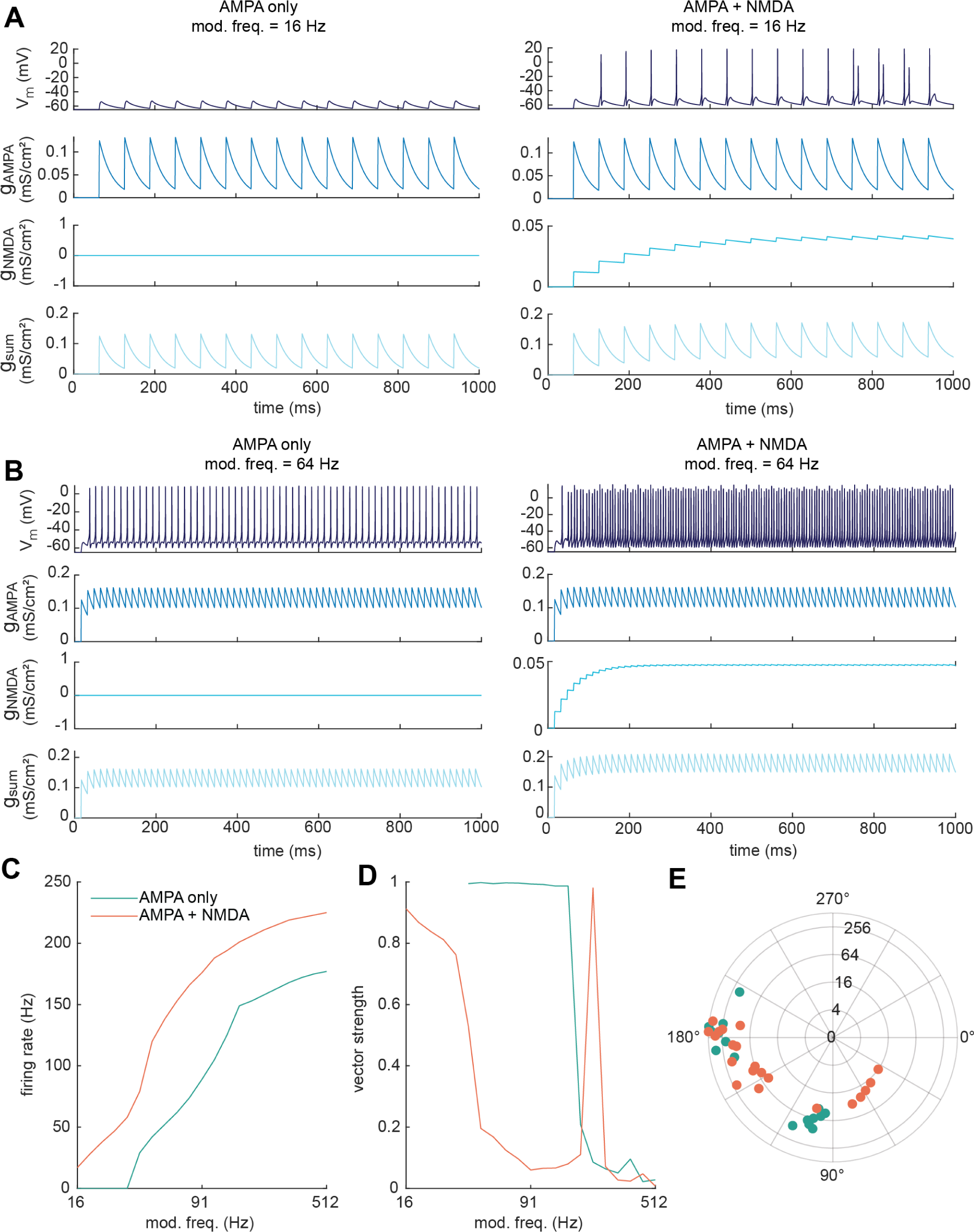
GluN2C/D-like NMDA conductance provides additive gain for rate coding in model IC neuron. ***A,*** Model neuron response to a 1 second, 16 Hz input train when the synapse contains an AMPA conductance (left panel) or both AMPA and NMDA conductances (right panel). To mimic GluN2C/D-containing NMDA receptors, the NMDA conductance was not voltage dependent. ***B,*** Model neuron response to a 1 second, 64 Hz input train when the synapse contains an AMPA conductance (left panel) or both AMPA and NMDA conductances (right panel). ***C,*** Rate modulation transfer functions for modulation frequencies ranging between 16 – 512 Hz at 4 steps/octave. Addition of the NMDA conductance (AMPA + NMDA, orange line) increased the firing rate at all modulation frequencies tested compared to the AMPA conductance alone condition (blue line). ***D,*** Temporal modulation transfer functions in response to the same modulation frequencies shown in ***C***. The model neuron exhibited strong phase locking over the mid-range of input frequencies in the AMPA only condition (blue line), and poorer phase locking in the AMPA plus NMDA condition (orange line). Note that the spike in phase locking at 215 Hz for the AMPA plus NMDA condition reflects instance where the sustained firing rate of the model neuron happened to align well with the input frequency. ***E,*** Distribution of firing phases for the AMPA conductance alone (blue dots) and the AMPA plus NMDA conductance (orange dots) conditions. Each dot represents a single modulation frequency. Concentric circles on the polar plot indicate modulation frequency.

### Membrane resistance alters modulation of rate coding by GluN2C/D-containing NMDARs

IC neurons are heterogeneous in their intrinsic physiology (reviewed in Drotos and Roberts, 2024), and these differences may alter how AMPA and NMDA receptors contribute to temporal and rate coding for stimuli. To examine how membrane resistance influences temporal integration, we simulated rate and temporal modulation transfer functions (MTFs) to input trains with frequencies between 16 – 512 Hz, as above, in model neurons where the leak conductance (*g_L_*) was set to 0.282, 0.50, 0.75, 1.99, and 5.3 mS/cm^2^ (corresponding to membrane resistances of 502, 283, 189, 71, and 27 MΩ, respectively, assuming 15 µm cell diameter). At the three lowest values for *g_L_*, neurons exhibited high-pass rate MTFs with a relatively smooth increase in firing rate as input frequency increased (**Figure 8A-C**, left panel). These cells also exhibited temporally precise firing at slow input frequencies, but the temporal precision of firing fell apart at higher input frequencies, and overall, the firing rate provided a more consistent indicator of input frequency. In addition, in these cells, the NMDA conductance contributed additive gain to the rate MTFs that shifted the input-output functions upwards along the *y*-axis, towards higher firing rates.

**Figure 8.**
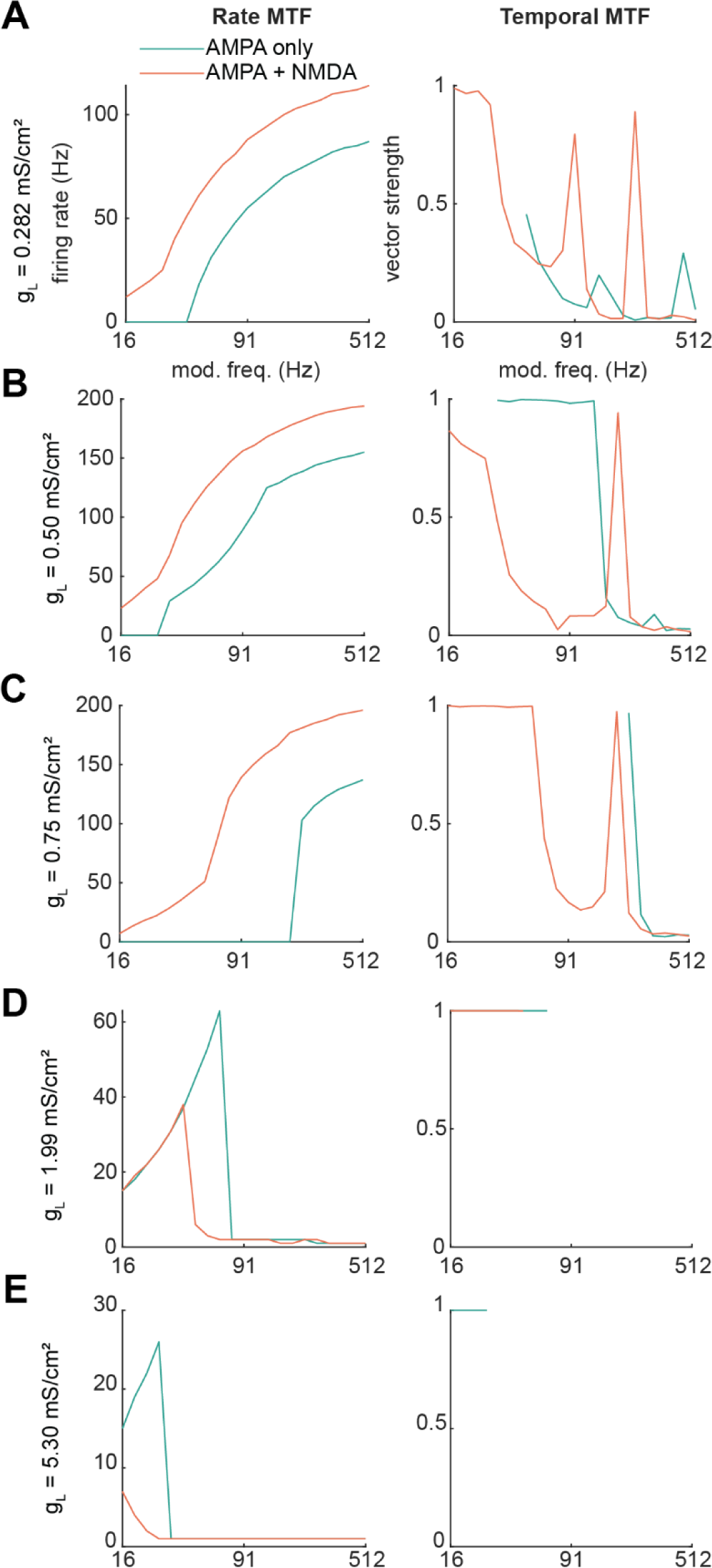
GluN2C/D-like NMDA conductance interacts with membrane resistance and AMPA conductance to shift temporal and rate coding for trains of inputs in the neuron model. Model neuron rate and temporal modulation transfer functions for leak conductances of *A,* 0.282, *B,* 0.50, *C,* 0.75, *D,* 1.99, and *E,* 5.30 mS/cm^2^. For a spherical neuron with a 15 µm diameter, these leak conductances correspond to 502, 283, 189, 71, and 27 MΩ membrane resistances, respectively. Blue lines indicate responses of the model neuron in the AMPA conductance alone condition, while orange lines indicate responses when the model synapse contained both AMPA and NMDA conductances.

As the leak conductance increased to 1.99, and 5.3 mS/cm^2^, we observed two major shifts: first, neurons increasingly exhibited better phase-locking and poorer rate coding of the input frequency, and second, addition of the NMDA conductance began to decrease, rather than increase, the gain of the rate MTF (**Figure 8D,E**). In our model, this was due to the additional conductance causing depolarization block, which shifted the response profile of these cells to an onset response. Note that rate MTFs in **Figure 8D,E** decreased to one spike per stimulus train, not zero, at higher input frequencies, and vector strengths were only calculated when input trains elicited >4 spikes per train. While it is likely that adding low-voltage-activated K^+^ conductances and/or other voltage-gated conductances to the model could convert onset firing neurons to phasic firing neurons, our main goal here was to mimic VIP neurons and other IC neurons with sustained firing, where such conductances are not prominent (Goyer et al., 2019). Overall, our modeling results highlight that the intrinsic physiology of a neuron influences both how that neuron uses temporal and rate codes for stimuli and how NMDA receptors shape these codes.

### GluN2C/D-containing receptors enhance gain for encoding AM stimuli

The results from our optogenetics experiments and neuron model suggest that NMDA receptors shape temporal and rate coding of synaptic stimuli. In the IC, temporal and rate codes are particularly important for encoding amplitude-modulated (AM) sounds, as IC neurons can encode the modulation frequency of AM stimuli with changes in firing rate and/or by phase-locking their firing to the modulation waveform (Krishna and Semple, 2000). Based on our model results, we hypothesized that GluN2C/D-containing NMDA receptors contribute to rate coding in the IC by broadening the time window for integrating phase-locked synaptic inputs and enhancing firing rates across different modulation frequencies. In support of this hypothesis, Zhang and Kelly showed that blocking NMDARs decreased rate coding in IC neurons while leaving temporal coding of AM stimuli intact (Zhang and Kelly, 2001), similar to what we observed in our model neuron (**Figures 7,8**). To examine how GluN2C/D-containing NMDARs influence rate coding for AM stimuli in the mouse IC, we made juxtacellular (loose-patch) current-clamp recordings from IC neurons in awake, headfixed mice while playing 1 s broadband noise bursts (4-64 kHz) that were amplitude modulated with modulation frequencies of 16 – 512 Hz and 100% modulation depths (**Figure 9A**).

**Figure 9.**
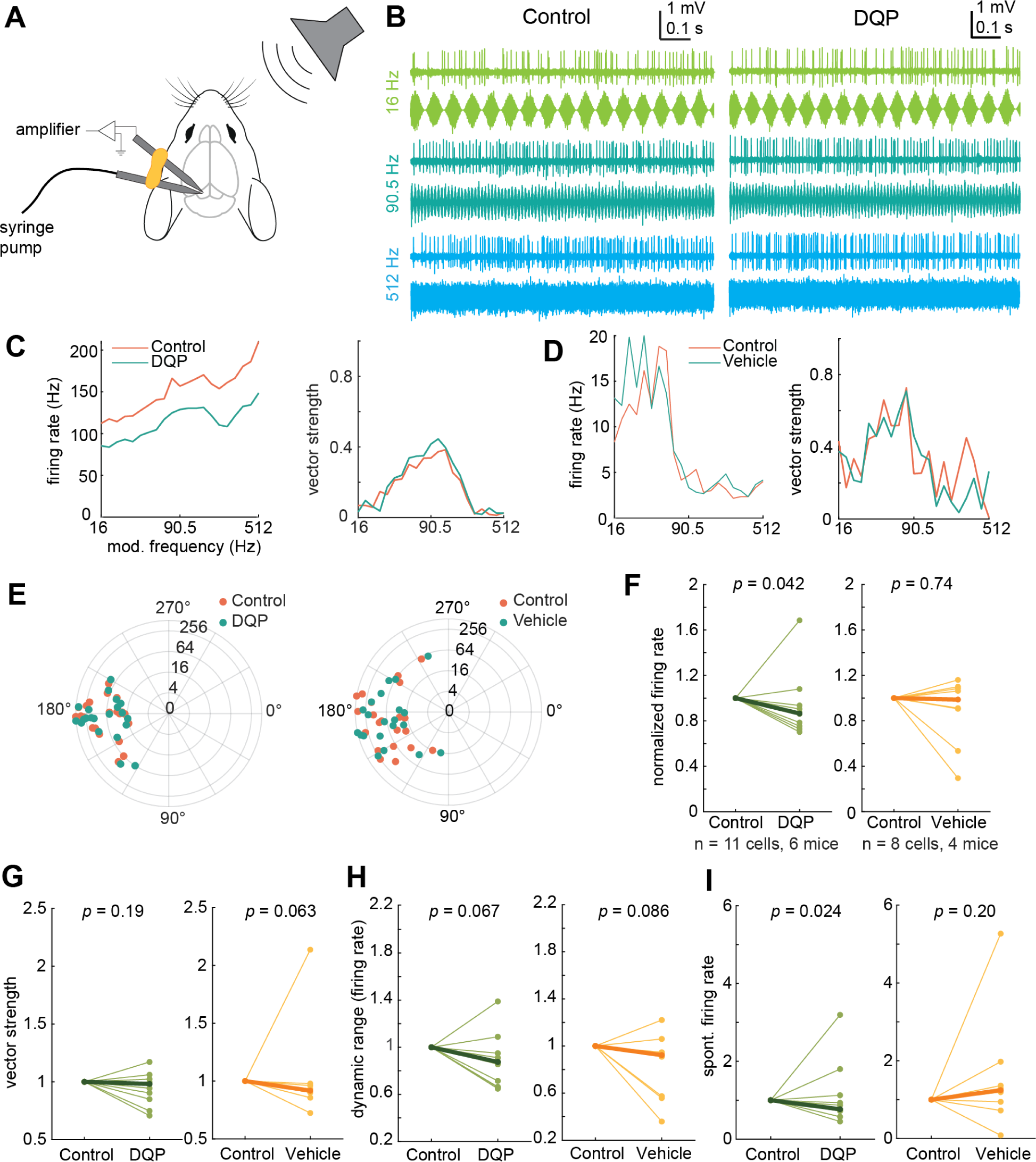
GluN2C/D-containing NMDARs enhance sound-evoked and spontaneous firing rates in vivo. ***A,*** Schematic of in vivo recording setup. Adapted from (Petrucco, 2020). In vivo juxtacellular recordings were performed in the IC of awake, headfixed mice using glass electrodes, and DQP or vehicle solution was infused using a piggyback pipette connected to a syringe pump. ***B,*** Juxtacellular recordings show both rate and temporal coding of the modulation frequency for AM noise bursts. In this example cell, the firing rate of the neuron increased with increasing modulation frequency. Second panel, the response of the same cell after infusion of the GluN2C/D-specific antagonist DQP. ***C,*** Blocking GluN2C/D-containing NMDA receptors via infusion of DQP decreased the firing rate of the example neuron from ***B*** at all modulation frequencies tested (blue line, left panel). Application of DQP had minimal or no effect on the vector strength of the response at any modulation frequency (right panel). ***D,*** Infusion of vehicle solution into the IC did not alter the firing rate (left panel) or vector strength (right panel) of the response to AM noise bursts. ***E,*** Infusion of DQP (left panel) or vehicle (right panel) did not alter the phases at which firing was phase locked in one DQP example neuron and one vehicle example neuron (orange is pre-infusion, blue is post-infusion). Each dot represents a single modulation frequency. Concentric circles on the polar plot indicate modulation frequency. ***F,*** DQP significantly decreased the number of spikes elicited during the sound stimulus (left panel, Z = 56, *p* = 0.042, median difference = -1.11 Hz, while vehicle infusion did not change the spike counts (right panel, Z = 21, *p* = 0.74, median difference = -0.062 Hz). ***G,*** DQP infusion (left panel) did not change the average vector strength of the response to AM stimuli (Z = 41, *p* = 0.19, median difference = -0.008), and vehicle infusion (right panel) also did not affect vector strength (Z = 15, *p* = 0.063, median difference = -0.018). ***H,*** Infusion of DQP (left panel) did not change the dynamic range of the firing rates over which IC neurons encoded AM modulation frequency (Z = 54, *p* = 0.067, median difference = -0.13 Hz), and this was also not altered by infusion of vehicle solution (right panel) (Z = 30.5, *p* = 0.086, median difference = -0.11 Hz). ***I,*** DQP decreased the overall spontaneous firing rate of IC neurons between stimulus presentations (left panel, Z = 58, *p* = 0.024, median difference = -3.03 Hz), while vehicle infusion had no effect on spontaneous firing rate (right panel, Z = 8, *p* = 0.20, median difference = 0.76 Hz).

In line with previous studies, we found that IC neurons encode AM stimuli via changes in firing rate (rate coding) and by phase locking their firing to the modulation waveform (temporal coding) (example cell response, **Figure 9B**). We next found that local infusion of 20 µM DQP, a GluN2C/D-selective antagonist, significantly decreased the firing rate during AM stimuli (signed-rank test, *p* = 0.042, median difference = -1.11 Hz) (**Figure 9C,F**) but had no effect on the average phase-locked responses (vector strength, signed-rank test, *p* = 0.19, median difference = -0.008) (**Figure 9C,G**) or the phase of the modulation frequency at which the neuron fired (**Figure 9E**). Overall, blocking GluN2C/D-containing receptors did not alter the shape of rate or temporal modulation transfer functions, nor did it significantly alter the relative change in firing rates (max – min) across the range of AM stimuli (dynamic range, *p* = 0.067, median difference = -0.13 Hz, **Figure 9H**). However, blocking GluN2C/D-containing receptors did significantly decrease the spontaneous firing rate during periods between stimulus presentations (*p* = 0.024, median difference = -3.03 Hz, **Figure 9I**). Together, these results suggest that GluN2C/D-containing receptors influence rate coding in IC neurons by providing a tonic, additive boost in gain to both spontaneous and sound-evoked synaptic inputs.

In addition, we ran control experiments to evaluate whether changes in firing could be due to infusion of solution into the IC or drift in the firing rate over the course of each recording. We found that infusion of vehicle solution did not alter the firing rate during AM stimuli (*p* = 0.74, median difference = -0.062 Hz), the vector strength of the response (*p* = 0.063, median difference = -0.018), the dynamic range in response to AM stimuli (*p* = 0.086, median difference = -0.12 Hz), or the spontaneous firing rate between sound presentations (*p* = 0.20, median difference = 0.76 Hz) (vehicle panels, **Figure 9D-I**).

## Discussion

In this study, we found that GluN2C/D-containing NMDARs shape synaptic integration and auditory processing in the IC. By combining optogenetics with whole-cell recordings, we found that NMDARs in VIP neurons are activated at resting potential by commissural projections. By using puffs of glutamate and whole-cell recordings combined with GluN2C/D-specific pharmacology, we showed that GluN2C/D-containing NMDARs activate at resting potential in VIP neurons. We demonstrated that 91% of VIP neurons express mRNA for the GluN2D subunit. We also found that *Glun2c* and *Glun2d* mRNA is prevalent throughout the IC, including in many non-VIP neurons, suggesting that GluN2C/D-containing NMDARs exert widespread influence over excitatory postsynaptic responses in the IC. Additionally, we demonstrated that VIP neurons can also express GluN2A/B-containing receptors, suggesting that different types of NMDARs may play distinct roles in these cells. Finally, we showed that GluN2C/D-containing receptors facilitate temporal summation in VIP neurons in vitro, provide additive gain to neuron input-output functions in silico, and enhance the spontaneous and sound-evoked firing rates of IC neurons in vivo. Thus, NMDAR diversity shapes synaptic integration and thus auditory processing in the IC.

### VIP neurons express NMDARs with GluN2D subunits

One of the most intriguing findings from our study was that the NMDAR currents elicited at resting potential in VIP neurons were predominately attributable to GluN2D-containing NMDARs rather than GluN2C. This result was unexpected for two reasons: first, in most brain regions, expression of *Glun2d* mRNA is developmentally regulated and disappears in adulthood (Monyer et al., 1994; Wenzel et al., 1996). Second, in adult animals, GluN2D-containing NMDARs are conventionally thought to be restricted to extrasynaptic locations, for example, in dorsal horn neurons (Momiyama, 2000) and Golgi cells of the cerebellum (Misra et al., 2000; Brickley et al., 2003), but there are some exceptions, such as in spinal cord laminar cells (Hildebrand et al., 2014). In addition, NMDARs are highly motile and can rapidly diffuse between extrasynaptic and synaptic sites (Tovar and Westbrook, 2002), as occurs during learning. During induction of NMDAR-LTP in the medial perforant path of the dentate gyrus, GluN2D-containing NMDARs are trafficked to the synapse where they contribute to synaptic transmission (Harney et al., 2008). Based on this, we hypothesize that GluN2D-containing receptors diffuse to synaptic locations on VIP neurons during learning. An alternative hypothesis is that GluN2D-containing NMDARs are natively found in the synapse and do not undergo a GluN2D-to-GluN2A/C switch during development.

Additionally, while we show here that many NMDARs on VIP neurons contain the GluN2D subunit, the full subunit composition of the receptor remains unknown. NMDARs are tetrameric receptors made up of four subunits, including two obligatory GluN1 subunits and a combination of GluN2A-D and/or GluN3 subunits (Traynelis et al., 2010). Our in situ hybridization results showed that 91% of VIP neurons express *Glun2d* mRNA. This could give rise to NMDARs that are diheteromeric with two GluN1 subunits and two GluN2D subunits. However, GluN1-GluN2D receptor assemblies have much slower kinetics than we observed in our study (decay τ = 10-100 ms), with decay time constants on the order of 2 seconds (Vicini et al., 1998). An alternative hypothesis is that GluN2D-containing receptors in VIP neurons are triheteromeric GluN1-GluN2B-GluN2D complexes. These receptors display properties that are intermediate to those of GluN1-GluN2B and GluN1-GluN2D receptors, including slightly faster kinetics than GluN1-GluN2D assemblies (Yi et al., 2019) and a reduced sensitivity to Mg^2+^ block compared to GluN1-GluN2B assemblies (Huang and Gibb, 2014). In addition, receptors containing GluN2B tend to be more motile than NMDARs with GluN2A subunits (Groc et al., 2006). Thus, this proposed GluN1-GluN2B-GluN2D assembly may also help explain how GluN2D-containing receptors contribute to synaptic transmission in mature VIP neurons. This conclusion is also supported by our CIQ experiments: we found a small effect of CIQ in the puff experiments and no effect of CIQ in the optogenetic activation experiments. The ability of CIQ to potentiate GluN2C/D-containing receptors is diminished in triheteromeric receptor assemblies compared to diheteromeric ones (Mullasseril et al., 2010), suggesting that the receptors studied here may be triheteromeric.

### Distribution of NMDARs on VIP neurons is synapse-specific

Previous work from our lab showed that NMDAR activation at resting potential on VIP neurons occurs with commissural inputs but not inputs from the DCN (Goyer et al., 2019). This suggests that the distribution of NMDARs on VIP neuron synapses is pathway-dependent, with commissural inputs targeting synapses that are enriched in GluN2D-containing receptors while DCN inputs target synapses predominated by AMPARs and possibly GluN2A/B-containing NMDARs. Heterogeneity in NMDAR composition at IC neuron synapses is also supported by the results from our in vitro and in vivo experiments, where some cells exhibited greater magnitude changes in response to blockade of GluN2C/D-containing NMDARs as compared to other cells. Such differences in receptor distribution could underlie functional differences between these synapses, as has been seen in other brain regions such as neocortex (Kumar and Huguenard, 2003). For example, commissural inputs may be integrated over longer time scales than those from the DCN, and a neuron may require more DCN inputs within a smaller time window to generate a postsynaptic spike.

NMDAR subtype distribution might also affect synaptic plasticity. GluN2A/B-containing NMDARs have been well-studied for their role in the initiation of long-term potentiation (LTP) (Paoletti et al., 2013). In the IC, previous studies show that electrical stimulation of the lateral lemniscus can induce LTP in IC neurons (Hosomi et al., 1995), and NMDARs are required for this process (Zhang and Wu, 2000; Wu et al., 2002). Our glutamate puff experiments showed that VIP neurons also express GluN2A/B-containing NMDARs, indicating that at least some VIP neuron synapses may exhibit AMPAR-dependent LTP (Malenka and Nicoll, 1993; Rebola et al., 2010; Hunt and Castillo, 2012). It will be important for future studies to determine the contributions of GluN2D trafficking and GluN2A/B expression to synaptic plasticity in the IC.

### GluN2D-containing receptors may alter rate coding for stimuli in the IC

Auditory structures primarily encode changes in sound amplitude envelopes in two ways: using temporal codes, where neurons phase-lock their firing to the modulation waveform, and using rate codes, where a neuron’s firing rate changes based on the frequency of the amplitude modulation (AM). While early auditory structures such as the cochlear nucleus primarily use temporal codes for AM stimuli (Rhode and Greenberg, 1994), neurons in auditory cortex primarily use rate codes (Yin et al., 2011). Since many IC neurons only phase lock to lower AM modulation frequencies (Rees and Møller, 1983; Rees and Palmer, 1989; Krishna and Semple, 2000) but exhibit strong and diverse dependencies of firing rate on changes in AM modulation frequency (Krishna and Semple, 2000; Nelson and Carney, 2007; Geis and Borst, 2009; Kim et al., 2020), the IC has been proposed as a critical site for the temporal-to-rate code transition that occurs between brainstem and cortex.

Our model and in vivo data highlight multiple important considerations for this transition. First, our model VIP neuron (**Figure 7**) and in vivo data (**Figure 9**) show that NMDA receptors contribute additive gain for rate coding of AM stimuli, providing a boost that pushes the neuron closer to spike threshold and increases the overall firing rate across modulation frequencies. This provides evidence for a mechanism that may promote rate coding in IC neurons by enhancing the dynamic range over which neurons encode AM modulation frequency using changes in firing rate. Second, when we investigated how adding an NMDA conductance altered temporal and rate coding in our model neuron, we found that the coding shifts were highly dependent on the input resistance of the neuron. Cells modeled with a lower leak conductance (Figure8A-C) and thus higher input resistance had clear rate coding for AM stimuli but exhibited poorer phase-locking. Addition of the NMDA conductance to the higher input resistance models provided additive gain which increased firing output, similar to what was seen in our in vivo experiments (**Figure 9C,F**). However, neurons modeled with a higher leak conductance and thus lower input resistance tended to exhibit stronger phase locking and poorer rate coding. In addition, adding the NMDA conductance to these models pushed the cell into depolarization block at higher input frequencies, thus turning the model neuron into a cell with an onset response profile.

Overall, our results show that GluN2C/D-containing NMDA receptors interact with AMPA receptors and passive membrane properties to shape rate and temporal coding in many IC neurons, helping give rise to the diverse ways that IC neurons encode AM stimuli and other temporally modulated sounds.

## Acknowledgements

The authors would like to thank Yoani Herrera, Marina Silveira, Pierre Apostolides, Marie Walicki, Will Birdsong, and Kelly Sovacool for helpful discussions and advice. This work was supported by a Rackham Graduate Student Research Grant (ACD), the National Science Foundation Graduate Research Fellowship Program under Grant No. DGE 1256260 (ACD), the National Institutes of Health Grant R01 DC018284 (MTR), and the National Institutes of Health Grant F31 DC021344 (ACD).

## Author contributions

ACD and MTR conceived of the study and designed experiments. ACD, VB, and RLZ conducted experiments. ACD analyzed data, prepared figures, and prepared the initial draft of the manuscript. ACD and MTR revised the manuscript, and all authors approved the submitted version.

